# Ribonucleotide incorporation characteristics around yeast autonomously replicating sequences reveal the labor division of replicative DNA polymerases

**DOI:** 10.1101/2020.08.27.270728

**Authors:** Penghao Xu, Francesca Storici

**Affiliations:** School of Biological Sciences, Georgia Institute of Technology, Atlanta, GA, 30332, USA

## Abstract

Ribonucleoside monophosphate (rNMP) incorporation in DNA is a natural and prominent phenomenon resulting in DNA structural change and genome instability. While DNA polymerases have different rNMP incorporation rates, little is known whether these enzymes incorporate rNMPs following specific sequence patterns. In this study, we analyzed a series of rNMP incorporation datasets, generated from three rNMP mapping techniques, and obtained from *Saccharomyces cerevisiae* cells expressing wild-type or mutant replicative DNA polymerase and ribonuclease H2 genes. We performed computational analyses of rNMP sites around early and late firing autonomously replicating sequences (ARS’s) of the yeast genome, from which bidirectional, leading and lagging DNA synthesis starts. We find the preference of rNMP incorporation on the leading strand in wild-type DNA polymerase yeast cells. The leading/lagging-strand ratio of rNMP incorporation changes dramatically within 500 nt from ARS’s, highlighting the Pol δ - Pol ε handoff during early leading-strand synthesis. Furthermore, the pattern of rNMP incorporation is markedly distinct between the leading the lagging strand. Overall, our results show the different counts and patterns of rNMP incorporation during DNA replication from ARS, which reflects the different labor of division and rNMP incorporation pattern of Pol δ and Pol ε.

## INTRODUCTION

Ribonucleoside monophosphates (rNMPs), the units of RNA are abundantly found incorporated in DNA from bacterial to mammalian cells (reviewed in (1)), leading to DNA structural change (2), genome instability (3) and disease (4–6). Yet, still much needs to be learned about rNMP patterns in DNA, and about the biological and clinical significance of rNMPs in genomic DNA. A proven, major cause for rNMP presence in DNA is misincorporation by replicative polymerases during the DNA replication process. It has been shown that abundant rNMPs are incorporated during DNA replication by the three replicative DNA polymerases of *Saccharomyces cerevisiae*, DNA Pol α, Pol δ, and Pol ε, which have different rNMP incorporation rates *in vitro* in physiological conditions (7, 8). On average, Pol α incorporates one rNMP per 625 nt, Pol δ one rNMP per 5,000 nt, and Pol ε one rNMP per 1,250 nt (8, 9). To distinguish an incoming deoxyribonucleotide triphosphate (dNTP) from a ribonucleotide triphosphate (rNTP), each of these replicative DNA polymerases (Pol α, Pol δ and Pol ε) contains a tyrosine in the active site functioning as a steric gate to block the rNTPs (10). Hence, point mutations at or next to this tyrosine residue in the catalytic subunit of replicative DNA polymerases (*pol1-Y869A* and *pol1-L868M* for Pol α, *pol2-M644G* for Pol ε, *pol3-L612M and pol3-L612G* for Pol δ) lead to a deficiency in the discrimination of dNTPs from rNTPs by the enzymes, and thus cause a higher level of rNMP incorporation during DNA replication (11, 12).

The principal repair mechanism of rNMP misincorporation is ribonucleotide excision repair (RER), which is induced by ribonuclease (RNase) H2 (13). As the first step of RER, RNase H2 recognizes rNMPs embedded in DNA and cleaves 5’ to the incorporated rNMPs. Hence, cells containing a null allele of the catalytic subunit of RNase H2 (*rnh201*-null for yeast and RNase H2A^-/-^ for mammalian cells) cannot perform the RER process and display a higher number of rNMP presence in their genomic DNA (7, 14).

Thanks to the recent development of the ribose-sequencing techniques, rNMP incorporation in DNA started to be studied in yeast *rnh201*-null cells carrying wild-type or mutant alleles of replicative polymerase α, δ and ε (11, 12, 15–17). These studies revealed the biased presence of rNMPs on the leading strand in wild-type DNA polymerase and in *pol2-M644G* cells, or on the lagging strand in *pol1-L868M, pol1-Y869A, pol3-L612M* and *pol3-L612G* mutant cells (11, 12, 16, 17). In *S. cerevisiae*, the DNA replication process with a leading and a lagging strand starts at autonomously replicating sequences (ARS’s) and proceeds bidirectionally requiring the activity of DNA polymerase α, δ and ε The lagging-strand synthesis is well known to occur in Okazaki fragments made by Pol α, followed by Pol δ. In contrast, whether leading-strand synthesis is catalyzed by Pol δ and/or Pol ε has been subject to ample debate (18, 19). Excitingly, recent experiments of replisome reconstitution *in vitro* using yeast purified proteins uncovered that an initial Okazaki fragment synthesized by Pol α and Pol δ on the lagging strand extends on the other side of the origin to prime leading strand synthesis by Pol ε (**Supplementary Figure 1**) (20, 21). Furthermore, successive studies conducted in yeast cells, exploiting leading/lagging biased distribution of rNMPs incorporated by a catalytic mutant of Pol α, δ or ε, have provided supportive evidence for the role of yeast DNA Pol δ in initiating leading strand DNA replication (12, 22). Nevertheless, the prevalence of the model for leading strand synthesis initiated by Pol δ and continued by Pol ε needs further confirmation, taking into consideration that the current work in yeast cells is mainly based on DNA polymerase mutants, which may not accurately represent what happens in cells carrying wild-type DNA polymerases.

**Figure 1.**
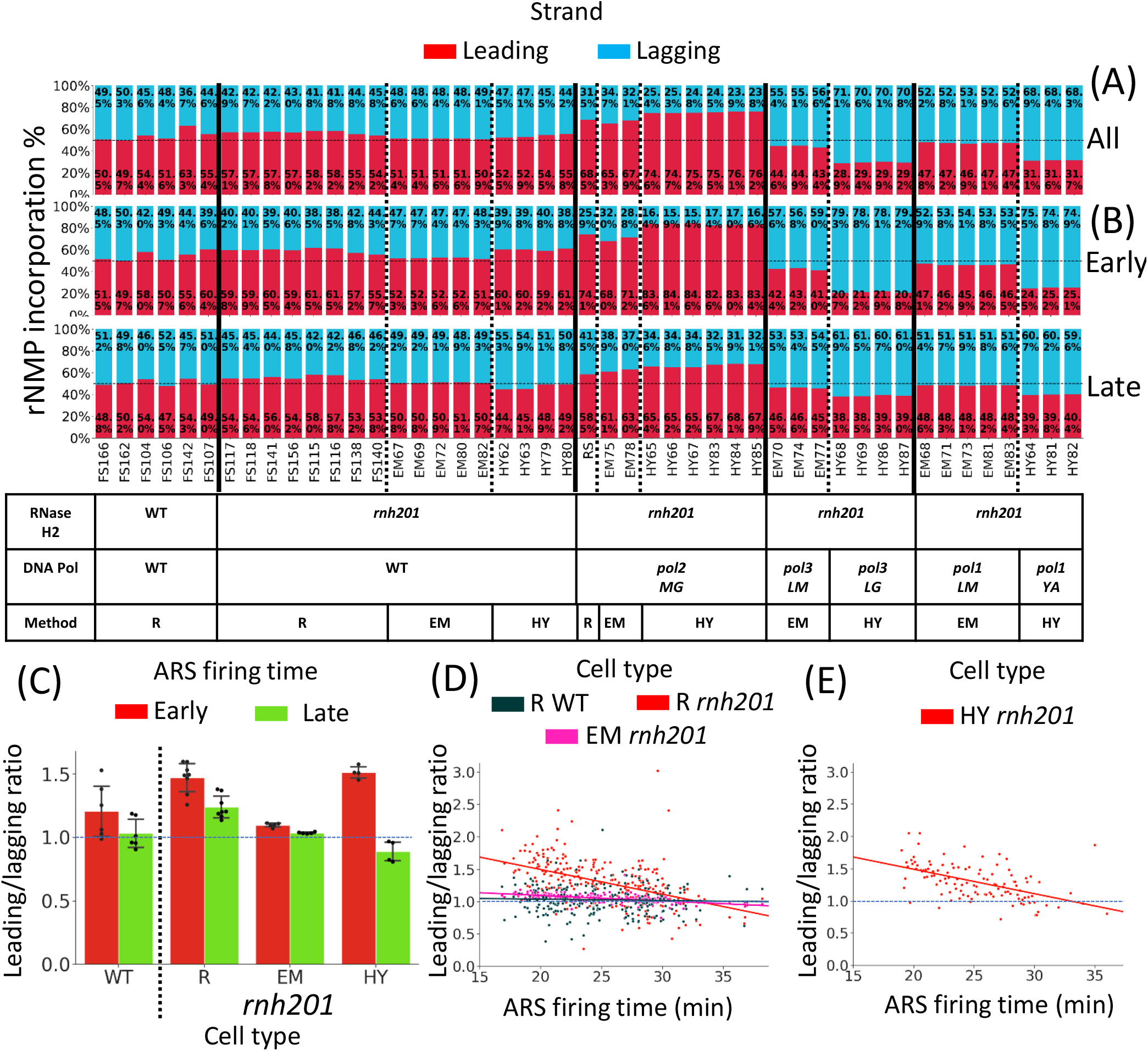
rNMP incorporation is prevalent on the leading strand in the presence of wild-type DNA polymerases α, δ and ε. **(A, B)** Bar graphs showing the % of rNMP incorporation on the leading (red bars) and lagging (blue bars) strands around all, early or late ARS’s, and the indicated rNMP libraries. The rNMP library name is indicated below each bar. ARS flank length = 15 kb. **(A)** Bar graph showing the percentage of rNMP incorporation for all confirmed ARS’s. All confirmed ARS’s in OriDB are used for ribose-seq and emRiboSeq libraries, while confirmed ARS’s in the L03 reference genome are used for RHII-HydEn-seq libraries (see Methods). **(B)** Bar graph showing the percentage of rNMP incorporation around early (top) and late (bottom) firing ARS’s. ARS’s with known firing time are divided into two halves based on their firing time. For ribose-seq and emRiboSeq libraries, firing time in Yabuki et al. 2002 (26) is used. Early-firing ARS’s have firing time < 24.7 min, n_early_ = 139, n_late_= 137. For RHII-HydEn-seq libraries, firing time in Zhou et al. 2019 (12) is used. Early-firing ARS’s have firing time < 26.7 min, n_early_ = 233, n_late_ = 232. The table below the bar graphs shows the genotypes of RNase H2 and DNA polymerases, as well as the technique used for the rNMP library preparation. R, ribose-seq libraries; EM, emRiboSeq libraries; HY, RHII-HydEn-seq libraries; *rnh201*, RNase H2 defective mutant; *pol2MG, pol2-M644G* mutant; *pol3LM:, pol3-L612M* mutant; *pol3LG, pol3-L612G* mutant; *pol1LM, pol1-L868M* mutant; *pol1YA, pol1-Y869A* mutant. **(C)** Bar graph showing the mean leading/lagging ratio of rNMP incorporation around early (red) and late (green) firing ARS’s in ribose-seq libraries with wild-type DNA polymerase and wild-type RNase H2, and RNase H2-defective ribose-seq, emRiboSeq, and RHII-HydEn-seq libraries. The thin dashed line marks a leading/lagging ratio = 1. **(D)** Scatter plot showing the relation between the leading/lagging ratio of rNMP incorporation around ARS’s with different firing times in the sacCer2 reference genome. The leading/lagging ratio around each ARS is calculated with maximum likelihood estimation. Clear decrease is found in *rnh201-null* ribose-seq libraries (Coefficient = −0.0382) and *rnh201-null* emRiboSeq libraries (Coefficient = −0.0085) while there’s only slight decrease in wild-type RNase H2 ribose-seq libraries (Coefficient = −0.0021). The thin dashed line marks a leading/lagging ratio = 1. **(E)** Scatter plot showing the relation between the leading/lagging ratio of rNMP incorporation around ARS’s with different firing times in the L03 reference genome. The leading/lagging ratio around each ARS is calculated with maximum likelihood estimation. Clear decrease is found in *rnh201-null* RHII-HydEn-seq libraries (Coefficient = −0.0380). The thin dashed line marks a leading/lagging ratio = 1.

Here, to provide new insights highlighting rNMP patterns and the division of labor at the replication fork around *S. cerevisiae* ARS sequences, we conducted a computational study using published datasets from three different rNMP mapping techniques, ribose-seq (16), emRiboSeq (11), and RHII-HydEn-Seq (12), and derived from cells of five different yeast strain backgrounds (BY4741, BY4742, YFP17, E134, and SNM106) expressing not only mutant but also wild-type polymerases, and containing RNase H2 wild-type or mutant alleles. The ribose-seq method uses alkali and the *Arabidopsis thaliana* tRNA Ligase (AtRNL) to directly capture rNMPs without capturing sites of nicks in DNA or Okazaki fragments(16, 23, 24). The emRiboSeq and RHII-HydEn-seq methods instead exploit either human RNase H2 or RNase HII from *Escherichia coli* combined with T4-quick ligase or T4 RNA ligase, to capture either the nicked sites right upstream of the rNMP or the nicked sites at the 5’ end of the rNMP with its downstream sequence, respectively (11, 12). We analyzed rNMP incorporation characteristics in specific regions around early and late firing *S. cerevisiae* ARS’s. We used either the OriDB database (25) with all 410 ARS’s from the reference genome SacCer2, 276 of which with known firing time are divided into two groups of early and late-firing ARS’s (26), or the 465 ARS regions identified in the L03 reference genome (12). We also divided the ARS’s of the L03 reference genome into two equal groups of early and late ARS’s. We find that rNMP distribution displays distinct leading/lagging-strand biases and patterns in different RNase H2 and DNA polymerase genotypes, which are also influenced by the ARS firing time. By studying the rNMP distribution and patterns at different distances from the ARS’s, we developed a model of rNMP incorporation rate on the leading and lagging strand, and we demonstrate the handoff from DNA Pol δ to Pol ε at the beginning of leading strand. Moreover, we identified unique patterns of rNMP incorporation on the leading and lagging strands, which reflect the rNMP incorporation preference of DNA polymerase Pol δ and Pol ε, and validating the handoff from DNA Pol δ to Pol ε at the beginning of the leading strand synthesis.

## MATERIALS AND METHODS

### Alignment of rNMP incorporation

All the libraries in the study are downloaded from NCBI. Library names in this study with corresponding SRR accession and BioProject accession are listed in **Supplementary Table 1**. For the ribose-seq and emRiboSeq libraries, the location and count of incorporated rNMPs are identified with Ribose-Map software on the SGD sacCer2 reference genome (27). For the RHII-HydEn-seq libraries, we downloaded the BigWig files aligning rNMP incorporation on the L03 reference genome. Then we converted them into BED files with bigWigToBedGraph and customized script (12, 28).

### ARS region information

For the ribose-seq and emRiboSeq, ARS annotations of OriDB were downloaded and transferred to the sacCer2 reference genome with Liftover software. Only confirmed ARS’s were included (n = 410) (29). Among them, 276 ARS’s have a known firing time (26). They were divided into two halves, early-firing ARS’s (T <= 24.7 min, n = 139) and late-firing ARS’s (T > 24.7 min, n = 137). For the RHII-HydEn-seq libraries, a total of 465 ARS’s with firing time were used in the study (12). They were also divided into two halves, early-firing ARS’s (T <= 26.7 min, n = 233) and late-firing ARS’s (T > 26.7 min, n = 232).

### Calculation of the flank of ARS’s and binning

The upstream and downstream 15-kb regions are considered as the flanks of an ARS. The 5’-upstream flank of ARS’s for both the Watson and Crick strands corresponds to the lagging strand, and the 3’-downstream flank of ARS’s corresponds to the leading strand. If two ARS’s are close to each other and the distance between them is smaller than 30 kb, the position of the replication fork collision is calculated with their firing times and the average fork moving speed of 1.6 kb/min (30). Their flanks are between the ARS’s location and the calculated collision position. We divided each ARS flank into 500-nt bins starting at each ARS location to analyze the variation of rNMP incorporation preference on the leading and lagging strands during the DNA replication process. Hence, each ARS flank has a maximum number of 30 bins. Moreover, 1-kb flank length and 100-nt bin size are used for the analysis of rNMP incorporation at the very beginning of DNA replication.

### The ratio of rNMP incorporation in the leading and lagging strands

Using the number of rNMPs incorporated on the 15-kb flanks of the leading and lagging strands around early and late ARS’s of wild-type RNase H2 and *rnh201*-null libraries prepared by three techniques, we calculated the mean value of the leading/lagging ratio as shown in the bar plot in **Figure 1**. Moreover, an estimated leading/lagging ratio for the line chart is obtained using libraries of the same genotype by the following assumption and calculation.

For the libraries i = 1,2,3, …, N with the same genotype, let *X*_*i*_ and *Y*_*i*_ denotes the number of rNMPs located in the leading and lagging strand in the ith library. Assuming *a* and *b* are the rNMP incorporation rates in the leading and lagging strand at each nucleotide, and *r*_*i*_ is the rate of which the incorporated rNMP could be captured and located. Letting *L* denote the length of the reference genome, we have that the number of rNMP sites follows the Poisson distribution.

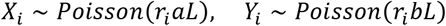

Then we can use Maximum likelihood estimation to calculate the rNMP incorporation rates *a* and *b*.

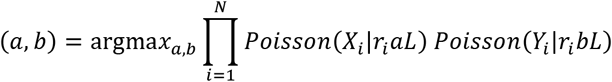

Solving the equation, we obtain,

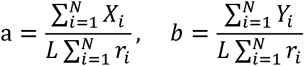

Hence, the estimated average leading/lagging ratio θ could be calculated.

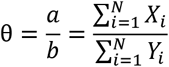

### Preference of rNMP incorporation

The heatmaps showing rNMP incorporation preference were generated as described in (24). Briefly, the four types of rNMPs were counted inside the 4-10-kb or 0-200-nt window of the ARS’s. The dinucleotides composed by the incorporated rNMP and its upstream neighbor (NR) or downstream neighbor (RN) were also counted. Each type of rNMP or dinucleotide was normalized based on the background dNMP frequency in the reference genome to obtain the normalized frequency. The background frequencies of the sacCer2 reference genome used by ribose-seq and emRiboSeq, and of the L03 reference genome used by RHII-HydEn-seq are listed in the **Supplementary Table 2**. The frequency was further normalized so that the sum of four types of rNMPs or four dinucleotides with the same rNMP in the same position is 1. Libraries with less than 100 rNMPs incorporated in the range of the ARS are excluded from the composition analysis, and those with less than 400 rNMPs incorporated around the ARS’s are excluded for the dinucleotide analysis to remove the large variation induced by a limited number of incorporated rNMPs. The box plots for dinucleotides (NR) around early and late-firing ARS’s with different genotypes were generated with 1.5 interquartile range (IQR).

### Statistical test

One-sided Mann-Whitney U test was performed to check whether the percentage of rNMPs incorporated on the leading strand was significantly higher or lower around early-firing ARS’s compared to late-firing ARS’s, and also to identify preferred rNMP incorporation patterns on the leading and lagging strands within 0-200-nt or within 4-10-kb windows around early and late-firing ARS’s.

## RESULTS

### rNMPs are preferentially incorporated on the leading strand of yeast genomic DNA derived from cells with wild-type DNA polymerases and wild-type RNase H2

To check the rNMP incorporation preference on the leading and lagging strands in yeast nuclear DNA, we calculated the percentage of the incorporated rNMPs in each strand separately within 15-kb flanks around each confirmed ARS in the *S. cerevisiae* genome for rNMP libraries prepared with three different rNMP mapping technique. We analyzed 15 ribose-seq libraries derived from 4 yeast strain backgrounds: E134, BY4741, BY4742, and YFP17 (16, 24), 15 emRiboSeq libraries derived from the Δl(−2)l-7B-YUNI300 background (11), and 17 RHII-HydEn-seq libraries derived from the Δl(−2)l-7B-YUNI300 background (12). The rNMP libraries with wild-type Pols were prepared by ribose-seq (N = 14), emRiboSeq (N = 5) or RHII-HydEn-Seq (N = 4) (11, 12, 24). Six of the 14 ribose-seq libraries of wild-type Pols are also RNase H2 wild-type. All other libraries of wild-type and mutant Pols are *rnh201*-null. Pol ε mutant *pol2-M644G* libraries were prepared by ribose-seq (N = 1), emRiboSeq (N = 2), or RHII-HydEn-Seq (N = 6) (11, 12, 16). Pol δ mutant *pol3-L612M* and *pol3-L612G* libraries were prepared by emRiboSeq (N = 4) or RHII-HydEn-Seq (N = 3), respectively (11, 12). Pol α mutant *pol1-L868M* and *pol1-Y869A* libraries were prepared by emRiboSeq (N = 5) or RHII-HydEn-Seq (N = 3), respectively (11, 12). All rNMP libraries used in the study, with their RNase H2 genotype, DNA polymerase genotype, strain name, and strain background, are listed in **Supplementary Table 1**. We utilized 410 confirmed OriDB ARS’s in the sacCer2 reference genome for ribose-seq and emRiboSeq libraries (25). For the RHII-HydEn-seq libraries, we instead used the 465 ARS’s with predicted firing time from the L03 reference genome described in Zhou et al. 2019 (12). To calculate the percentage of rNMPs present on the leading and lagging strand of all these libraries, we examined rNMPs within 15-kb flanks of the ARS’s, which cover the initiation and elongation tract of the leading and lagging strands. The termination zones of either leading or lagging strand synthesis are not included within these 15-kb flanks of the ARS’s (12). The percentage of rNMP incorporation on the leading and lagging strands in the 15-kb flanks around all confirmed ARS’s are distinct (**Figure 1A**). There are more rNMPs incorporated on the leading than on the lagging strand in all wild-type DNA polymerase libraries prepared by the three different techniques. This preference is more stable in *rnh201*-null libraries (17 out of 17), likely because the *rnh201* libraries have a larger number of incorporated rNMPs (11, 12, 24). However, we find that there is a preference for rNMP incorporation on the leading strand also in wild-type RNase H2 cells (5 out of 6, and 1 library shows no preference). These results underscore a prevalent and overall higher level of rNMP incorporation by DNA polymerases working on the leading strand.

Low fidelity mutants of DNA polymerases have a higher probability of misincorporating rNMPs in a strand-biased manner (9). As expected, our analysis finds a stronger preference of rNMPs on the leading strand of *pol2-M644G* mutant libraries, whose mutant is embedded in DNA Pol ε. Oppositely, there is a stronger preference for rNMP incorporation on the lagging strand of *pol1* and *pol3* mutant libraries (*pol1-L868M* and *pol3-L612M* for emRiboSeq libraries, *pol1-Y869A* and *pol3-L612G* for RHII-HydEn-seq libraries), whose mutants are in DNA Pol α and Pol δ (**Figure 1A**). We note that results obtained using RHII-HydEn-seq libraries show stronger strand bias in the mutant Pols. This could be due to the choice of mutants *pol1-Y869A* and *pol3-L612G*, which have higher rNMP incorporation frequency compared to *pol1-L868M*, vs. *pol3-L612M* used in the emRiboSeq libraries (11, 12), as well as the fact that the L03 reference genome was specifically generated from the Δl(−2)l-7B-YUNI300 background used for the preparation of these RHII-HydEn-seq libraries, or both cases may apply.

### The strand preference for rNMP incorporation is stronger around ARS’s with an early firing time

The different firing times of ARS’s affect the length of the synthesized leading and lagging strand tracts from the ARS’s. Usually, the replicated DNA sequence from an early-firing ARS tends to be longer than the replicated DNA sequence from a late-firing ARS before the ARS’s are colliding with each other. We wondered whether the firing time of the ARS’s would affect the leading/lagging-strand preference of rNMPs that we observed for wild-type and RNase H2 mutant cells of wild-type and mutant DNA polymerases. To address this question, we divided the ARS’s with known firing times into two halves based on their early or late firing time, as described in the Methods section. We then calculated the percentage of rNMP incorporation on the leading and lagging strands for the two groups with all rNMP libraries. In the RNase H2 wild-type and *rnh201*-null of wild-type DNA polymerase libraries, the rNMP preference is on the leading strand, and such preference weakens within the late-firing ARS’s compared to the early-firing ARS’s (*P* = 0.046, *N* = 6 for RNase H2 wild-type, and *P* = 3.2 × 10^−4^, *N* = 17 for *rnh201*-null libraries, respectively; **Figure 1B**). The weakened rNMP preference of the leading strand is also observed in the *pol2-M644G* mutant libraries for the late ARS’s compared to the early ARS’s (*P* = 2.9 × 10^−4^, *N* = 9, **Figure 1B**). As for the *pol1* and *pol3* mutant libraries (*pol1-L868M* and *pol3-L612M* for emRiboSeq libraries, *pol1-Y869A* and *pol3-L612G* for RHII-HydEn-seq libraries), in which the rNMPs are preferentially incorporated on the lagging strand, such preference is also less significant around the late-firing ARS’s compared to the early-firing ARS’s (*pol1*: *P* = 0.042, *N* = 8; *pol3*: *P* = 0.063, N = 7, **Figure 1B**).

We then performed the quantitative analysis by calculating the leading/lagging ratio of rNMP incorporation and obtain the mean value to determine the effect of ARS firing time on the preference of rNMP incorporation in wild-type DNA polymerase libraries. Results show a reduced leading/lagging ratio around the late-firing ARS’s in both RNase H2 wild type and *rnh201*-null ribose-seq libraries, which suggests that the leading-strand preference is weakened around late-firing ARS’s. The *rnh201*-null emRiboSeq libraries show a similar but more slight decrease, and *rnh201*-null RHII-HydEn-seq libraries show a stronger decrease **(Figure 1C**). To quantitatively determine how the increased firing time reduces the rNMP-incorporation preference on the leading vs. the lagging strand, we calculated the maximum likelihood estimation of the average leading/lagging ratio in each ARS flank, and we performed linear regression analysis with the ratios and the firing time for the ARS’s. Despite some variance due to the small number or rNMP incorporation in a single ARS flank, we observed a clear decrease of the leading/lagging ratio with the increased firing time in *rnh201*-null ribose-seq libraries (**Figure 1D**, *Coefficient* = −0.0382) and RHII-HydEn-seq libraries (**Figure 1E**, *Coefficient =* −0.0380). The decrease of the leading/lagging ratio is slighter in *rnh201*-null emRiboSeq libraries (**Figure 1D**, *Coefficient* = −0.0085) but is still stronger than wild-type RNase H2 ribose-seq libraries (**Figure 1D**, *Coefficient* = −0.0021). In summary, we found that both the leading-strand preference for rNMP incorporation in wild-type DNA Pols and *pol2-M644G* libraries and the lagging-strand preference in *pol1-L868M, pol1-Y869A, pol3-L612M*, and *pol3-L612G* libraries are reduced with increasing firing time. And the reduction is stronger with RNase H2-deficient libraries compared to wild-type RNase H2 libraries.

### DNA polymerase switches shape the rNMP preferences on the leading and lagging strands

During DNA replication in budding yeast, the leading and lagging strands are synthesized with DNA Pols α, δ, and ε. According to the recent *in vitro* studies and work in yeast with mutant Pols, the synthesis of the leading strand in yeast DNA starts with an extended Okazaki fragment (∼180 nt) made by DNA Pol δ, and DNA Pol ε takes the main part afterward (12, 20–22) (**Supplementary Figure 1**). Finally, DNA Pol ε switches back to DNA Pol δ when meeting the incoming lagging strand (12). The lagging strand is composed of short Okazaki fragments synthesized by DNA Pol α and Pol δ. Hence, the rNMP incorporation rate is expected to change during DNA replication on both the leading and lagging strands consistently following the division of labor of these Pols. Specifically, the rNMP incorporation rate on the leading strand would follow a high-low-medium pattern from the starting sites. However, the starting site of leading strand synthesis is upstream of the ARS (21). If we track the rNMP incorporation rate from the ARS, we should start with the Pol δ phase and in a low-medium pattern (**Figure 2A**). On the lagging strand, the rNMP incorporation rate would change with a periodical pattern of Okazaki fragment length (**Figure 2B**). Furthermore, the average rNMP incorporation rate change around combined ARS’s would be smoothed because there are always deviations in the position of the actual ARS’s relative to the annotated ones. With increasing distance from the ARS’s, on the leading strand, the rNMP incorporation rate should increase in wild-type DNA polymerase and *pol2-M644G* libraries and decrease in *pol3-L612M* and *pol3-L612G* libraries (**Figure 2C**). Since the deviation is usually larger than the length of the Okazaki fragment, the rNMP incorporation rate on the lagging strand should be steady as a horizontal line with small waves (**Figure 2D**). As a validation, we calculated the average value of rNMP per base (RPB) of wild-type DNA polymerases in *rnh201*-null ribose-seq libraries, which is the direct measurement of the rNMP incorporation rate. On the leading strand, the RPB gradually increases in the first 2.5 kb around early-firing ARS’s and in the first 4 kb around late-firing ARS’s (**Figure 2E**). On the lagging strand, the RPB keeps steady in the DNA replication process. Only small variations are found around early or late-firing ARS’s (**Figure 2F**). These results suggest that the trends of the rNMP incorporation rate during DNA replication are in line with our theoretical model shown in **Figure 2C** and **2D**.

**Figure 2.**
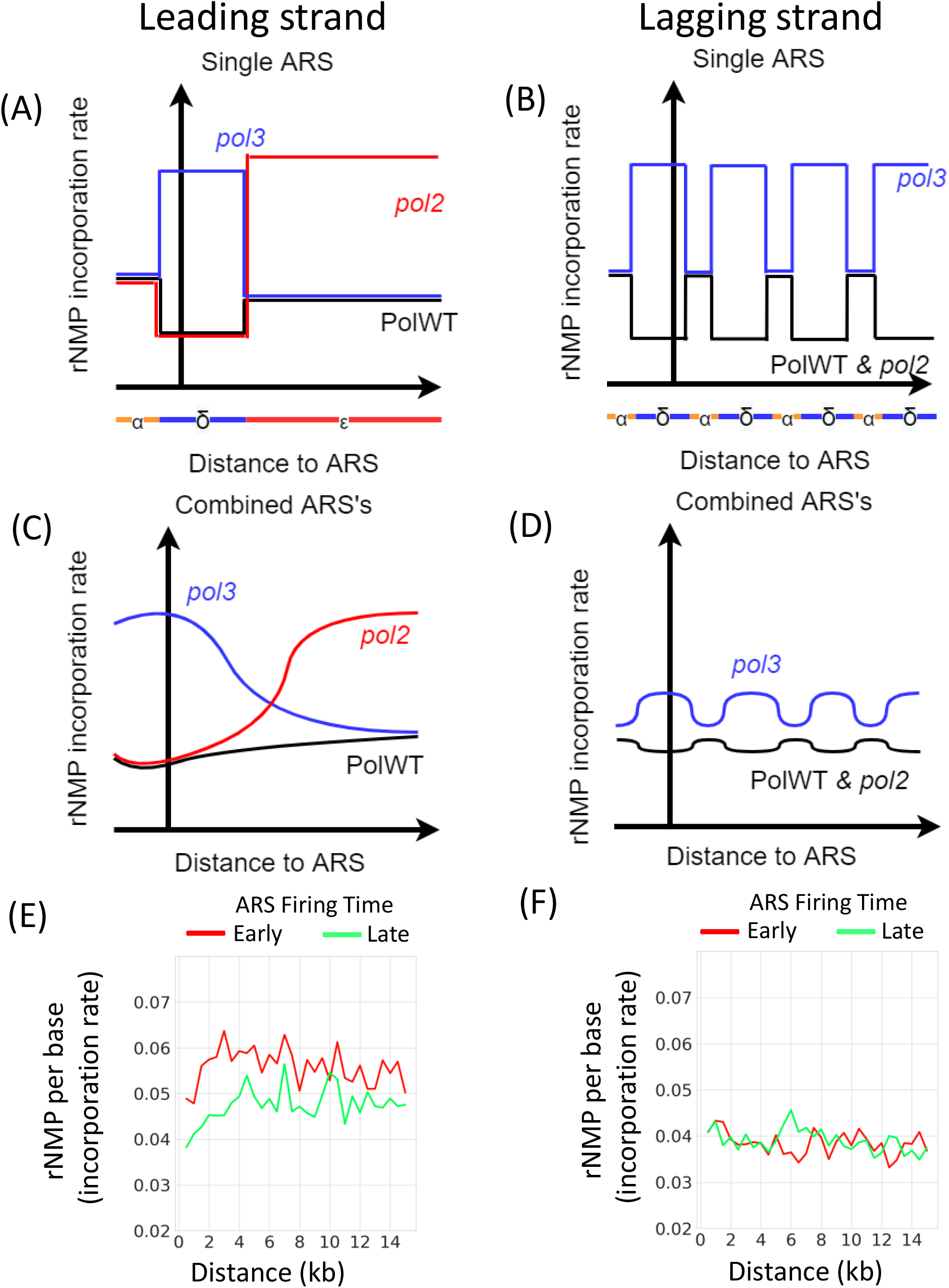
Model for the rate change of rNMP incorporation during DNA replication. **(A)** The rNMP incorporation rate on one leading strand of an ARS. The colored line below the X-axis shows the model of division of labor by DNA polymerases in Aria et al. 2019 and Zhou et al. 2019 (12, 21), in which the leading strand synthesis begins with a “lagging strand” primer initiated by DNA Pol α and extended by Pol δ. The leading strand synthesis begins on the upstream of the ARS location (Y-axis), which is shown as the leftmost of the X-axis. The rNMP-incorporation rate will change immediately when DNA replication is switched to a different DNA polymerase. Hence, there are 3 phases of the rate induced by DNA Pol α, Pol δ, and Pol ε synthesis. Only two phases (Pol δ, and Pol ε) can be found on the leading strand from the ARS location. **(B)** The rNMP incorporation rate on one lagging strand of an ARS. The rNMP-incorporation rate changes like a rectangular wave with the size of the Okazaki fragment. **(C)** The average rNMP-incorporation rate on the leading strands of combined ARS’s. When multiple ARS’s are considered, the rate is smoothed due to the deviation of the annotated ARS location. **(D)** The average rNMP-incorporation rate on the lagging strand of multiple ARS’s. The rate shows like a horizontal line with small waves. **(E)** Example of an average rNMP-incorporation rate on the leading strand. All ribose-seq *rnh201-*null libraries with wild-type DNA polymerases are used (N = 8). The increase of the rNMP incorporation rate at the beginning is in consensus with the pattern of wild-type DNA polymerase of the leading strand (black curve in **C**). **(F)** Example of an average rNMP incorporation rate on the lagging strand. All ribose-seq *rnh201-*null libraries with wild-type DNA polymerases are used (N = 8). There are only small variations in the ratio, which follows the pattern of wild-type DNA polymerase of the lagging strand (black curve in **D**).

Moreover, the leading/lagging ratio of rNMP incorporation has the same trend as the leading strand since the rNMP-incorporation rate on the lagging strand remains steady. And it is a better measurement of rNMP incorporation preference since it is insensitive to rNMP counts compared to the leading-strand and lagging-strand incorporation rate (**Supplementary Figure 2**). In wild-type DNA polymerase *rnh201-*null libraries, the leading/lagging ratio of rNMP incorporation increases at the beginning both around early and late-firing ARS’s (**Figure 3**). Since wild-type DNA Pol δ has a lower rNMP incorporation rate than wild-type Pol ε, the observed increase suggests that the working polymerase for DNA replication is gradually changing from DNA Pol δ to Pol ε. A similar increase is also evident in the wild-type RNase H2 libraries. The increase is weaker since the RER mechanism balances the leading strand preference, and there are more prominent variations after the increase due to the limited number of incorporated rNMPs. A similar but stronger increase occurs in RNase H2 deficient *pol2-M644G* mutant libraries since mutant DNA Pol ε has an even higher rNMP incorporation rate than wild-type Pol ε (**Figure 3**). In *pol3-L612M* RHII-HydEn-seq and *pol3-L612G* emRiboSeq libraries, the leading/lagging ratio of rNMP incorporation decreases at the beginning, both for early and late firing ARS’s. This is because mutant Pol δ has a higher rNMP incorporation rate than wild-type Pol ε (**Figure 3**). At the beginning of replication, all libraries have a leading/lagging ratio near 1 compared to the following windows since Pol δ is present on both leading and lagging strands. Overall, these analyses demonstrate that from the ARS center on the leading strand, Pol δ starts DNA synthesis with a handoff to Pol ε.

**Figure 3.**
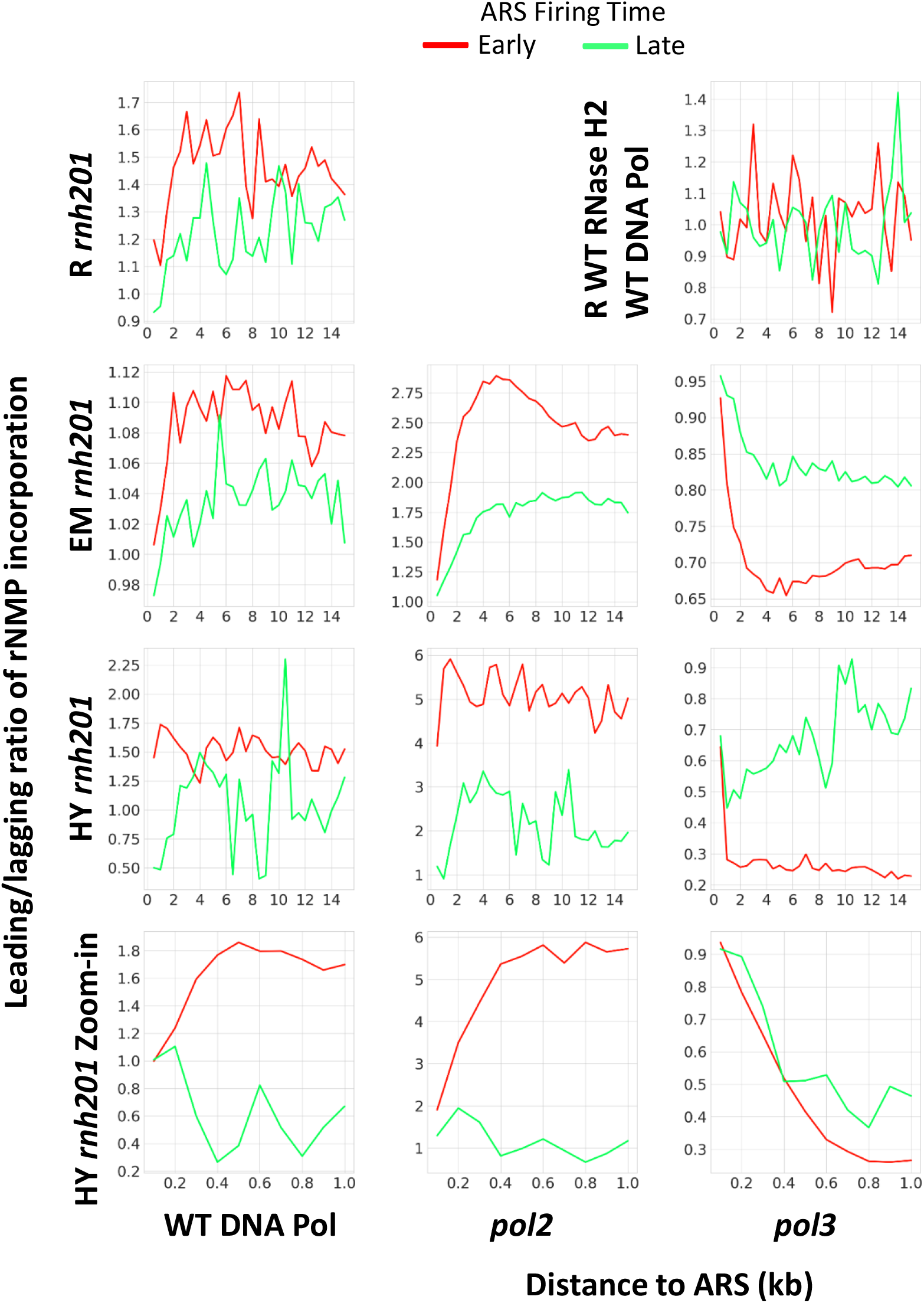
The leading/lagging ratio of rNMP incorporation changes during DNA replication. Maximum likelihood estimation is used to calculate the leading/lagging ratio in RNase H2 wild-type and wild-type DNA polymerase libraries, or *rnh201-null* wild-type DNA polymerase or mutant polymerase of ribose-seq, emRiboSeq or RHII-HydEn-seq libraries. ARS flank length = 15 kb and bin size = 0.5 kb are used for the plots, except for the RHII-HydEn-seq zoom-in plots in the last row, in which ARS flank length = 1 kb and bin size = 0.1 kb are used. R, ribose-seq libraries; EM, emRiboSeq libraries; HY, RHII-HydEn-seq libraries; *rnh201*, RNase H2 defective mutant; *pol2, pol2-M644G* mutant; *pol3, pol3-L612G* mutant for RHII-HydEn-seq libraries and *pol3-L612M* mutant for emRiboSeq libraries.

Furthermore, the length of the changing phase (increase or decrease) at the beginning of DNA replication also reflects when the switch in DNA polymerase occurs. For example, in wild-type DNA polymerase libraries of *rnh201*-null cells, at the beginning of DNA replication when DNA Pol δ takes all synthesis work on both the lagging and the leading strands, the rNMP incorporation rate is the lowest. Differently, when DNA Pol ε takes the synthesis work on the leading strand, the rNMP incorporation rate is the highest. Thus, the length of the increase phase marks the range of the occurring handoff from Pol δ to Pol ε. We find that the changing phase is 4 kb in mutant Pol ε (*pol2-M644G*) of emRiboSeq libraries and 1 kb in the same Pol ε mutant of RHII-HydEn-seq libraries, which is the handoff range from DNA Pol δ to Pol ε (**Figure 3**). The handoff range is longer than the 180-bp Pol δ tract in the Zhou et al. study (12). It is because in the Zhou et al. study the Pol δ tract refers to a selected range of nucleotides synthesized by Pol δ for the majority of ARS’s. Such selected range matches the range size of the most extreme leading/lagging ratio in our study, which corresponds to the beginning of the changing phase. The more rapid the increase or decrease of the leading/lagging ratio is at a certain distance from the ARS’s in the plots, the more frequently the polymerase handoffs happen at that distance around all ARS’s. Our analysis shows that the most rapid phase change occurs within the first 2 kb in the emRiboSeq libraries (**Figure 3**), and within the first 500 nt in the RHII-HydEn-seq libraries (**Figure 3**, zoom-in). Thus, the majority of ARS’s have completed the handoff in this phase. In addition, in the wild-type DNA polymerase libraries of *rnh201-*null cells prepared using any of the three different techniques, the length of the increasing phase around the late-firing ARS’s is always longer than that around early-firing ARS’s (**Figure 3**). This suggests that Pol δ usually synthesizes a longer tract before its handoff to Pol ε around late-firing ARS’s. This elongated increasing phase around late-firing ARS’s is less prominent in the Pol ε and Pol δ mutant libraries, suggesting that the length of the Pol δ-synthesized tract before the handoff to Pol ε can be affected by the firing time only in the wild-type DNA polymerase libraries. Moreover, in the wild-type DNA polymerase and *pol2-M644G* mutant RHII-HydEn-seq libraries, the leading/lagging ratio shows a slight decrease at the beginning around late-firing ARS’s, which may be related to Pol α to Pol δ switch.

In general, RHII-HydEn-seq libraries have a shorter changing phase compared to ribose-seq and emRiboSeq libraries, which likely suggests that the ARS annotation for the L03 reference genome has the least deviation. In addition, we find that in wild-type DNA Pol libraries, the *rnh201*-null ribose-seq, and emRiboSeq libraries have a similar magnitude of leading/lagging-ratio increase or decrease, while there is a much larger magnitude in RHII-HydEn-seq libraries (**Supplementary Figure 3**). The magnitude of increase and decrease is caused by different rNMP incorporation rates of the DNA polymerases. Hence, there is a higher rNMP incorporation rate in wild-type Pol ε, mutant Pol ε, and mutant Pol δ for the strain background of RHII-HydEn-seq libraries compared to the strain backgrounds of ribose-seq and emRiboSeq libraries.

### The rNMP composition on the leading and lagging strands is the same

We picked two windows for each ARS flanks to check the rNMP incorporation patterns in the leading and lagging strand. The first is a 4-10-kb window. In this window, the leading strand is mainly synthesized by DNA Pol ε because the handoffs from Pol δ to Pol ε around almost all ARS’s are completed at such distance from the ARS’s. The other window is the 0-200 nt, which corresponds to the most rapidly changing phase for all the three rNMP-mapping techniques, in which the leading strand is synthesized by both DNA Pol δ and Pol ε. For the 4-10-kb window, we analyzed 39 libraries prepared using the ribose-seq, emRiboSeq, or RHII-HydEn-seq techniques and derived from 5 different yeast strain backgrounds and 11 different genotypes (**Figure 4**). In our recent study by Balachander, Gombolay, Yang, Xu et al., 2020 (24), we revealed both conservations in rNMP composition, including low rU in *rnh201*-null strains in nuclear DNA, as well some variation across different strains of the same *S. cerevisiae* species. Notably, some strains had a very high frequency of rC, while both rC and rG had high frequency in other strains (24). Similar to the full double-stranded nuclear genome (24) in *rnh201*-null cells, rU is always the least abundant rNMP incorporated on both the leading and lagging strands around early as well as late ARS’s, independently from the genotype of the DNA polymerases (**Figure 4A, B**, and **Supplementary Figure 4A, B**). And rG is the least incorporated rNMP in wild-type RNase H2 cells on both the leading and the lagging strands around early as well as late ARS’s (**Figure 4A** and **Supplementary Figure 4A**). Moreover, there is no significant difference across the various polymerase alleles in libraries prepared by the three techniques, respectively (**Figure 4B, Supplementary Figure 4B**). We note that for the ribose-seq and RHII-HydEn-seq libraries of *rnh201*-null cells, rC is the most abundant rNMP, while both rC and rG are abundant in emRiboSeq libraries. This difference most likely reflects variation across the different strain backgrounds, as noted before (24). Here, we found that the composition of incorporated rNMPs is consistently similar between the leading and lagging strands around both early (**Figure 4A**) and late firing ARS’s (**Supplementary Figure 4A**) in all genotypes and libraries examined within the 4-10-kb window around the ARS’s. For the 0-200-nt window, there are fewer rNMPs incorporated, and libraries with less than 100 rNMPs incorporated in this window were excluded from the analysis. Among the 34 libraries examined, we find that the rNMP composition in the 0-200-nt window is similar to that in the 4-10-kb window. And no difference is found between leading and lagging strands and around early or late-firing ARS window (**Figure 4C, D**, and **Supplementary Figure 4C, D**). These results show that not only there is no difference between the rNMP composition on the leading vs. the lagging strand within both the 0-200 nt and the 4-10-kb windows, but the composition on these individual strands is also similar to the that found in the whole nuclear DNA of the same ribose-seq libraries of wild-type RNase H2 and *rnh201*-null cells and for the emRiboSeq libraries of *rnh201*-null cells studied in (24).

**Figure 4.**
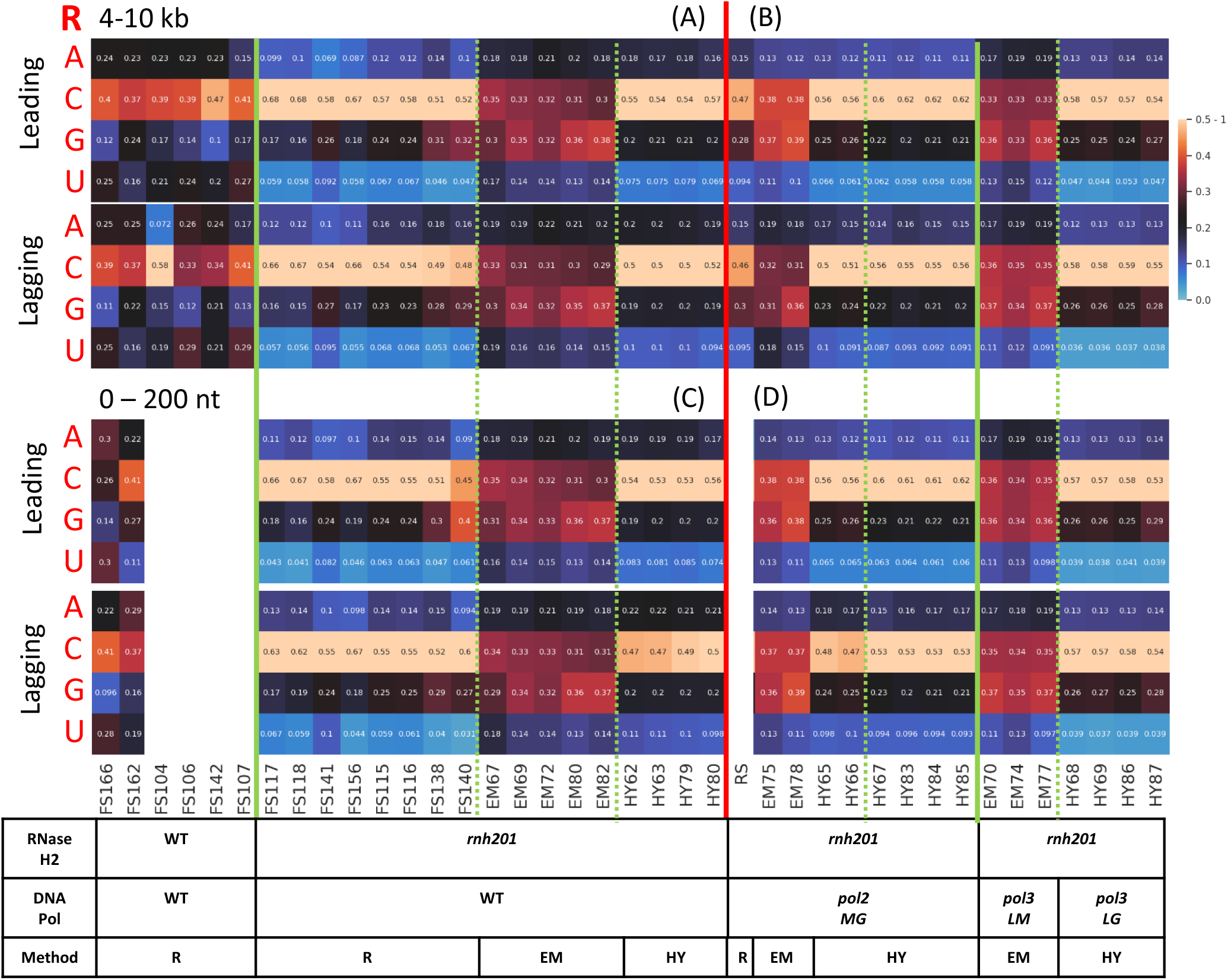
The composition of incorporated rNMPs around early-firing ARS’s on the leading and lagging strand is the same. Heatmap analyses with the normalized frequency of each type of rNMP (R: rA, rC, rG, or rU). The counts for each type of incorporated rNMP are normalized to the nucleotide frequencies of the 4-10 kb (top) and 0-200 nt (bottom) windows for the leading or lagging strand around early ARS’s from the sacCer2 reference genome for all the ribose-seq and emRiboSeq libraries, and from the L03 reference genome for all the RHII-HydEn-seq libraries. The sum of 4 types of rNMP frequency is further normalized to 1. Hence, 0.25 is the expected normalized frequency if there is no rNMP incorporation preference. The corresponding formula used is shown in the Methods section. The background nucleotide frequencies of ribose-seq and emRiboSeq (according to the sacCer2 reference genome), and RHII-HydEn-seq (according to the L03 reference genome) libraries are reported in **Supplementary Table 2A** and **2B**, respectively. Each column of the heatmap shows the results of a specific library. Some libraries with less than 100 rNMPs incorporated were excluded to generate the 0-200-nt window plots. The table underneath the heatmap shows the genotypes of RNase H2 and DNA polymerases, as well as the technique used for the rNMP library preparation. The thick, vertical, red line separates data obtained from wild-type DNA polymerase (**A** for 4-10 kb, **C** for 0-200 nt) from data obtained with mutant DNA polymerases (**B** for 4-10 kb, **D** for 0-200 nt). The thick, vertical, green lines separate data obtained from wild-type RNase H2 from those obtained with *rnh201-null* libraries of wild-type DNA polymerase, and data obtained from different mutant DNA polymerases of *rnh201*-null libraries. The dashed, green lines separate data obtained using different rNMP mapping techniques. Each row shows results obtained for a type of rNMP. The bar to the right shows how normalized frequencies are represented as different colors: black for 0.25; black to yellow for 0.25 to 0.5 – 1, and black to light blue for 0.25 to 0. R, ribose-seq libraries; EM, emRiboSeq libraries; HY, RHII-HydEn-seq libraries; *rnh201*, RNase H2 defective mutant; *pol2MG, pol2-M644G* mutant; *pol3LM, pol3-L612M* mutant for emRiboSeq libraries; *pol3LG, pol3-L612G* mutant for RHII-HydEn-seq libraries.

### The rNMP incorporation patterns on the leading and lagging strands are distinct

In contrast to the rNMP composition, we uncover that the sequence context of incorporated rNMPs is different on the leading and lagging strands. For the incorporated rNMPs on the leading and lagging strands within the same 4-10-kb window from the ARS’s, we determined the frequency in which each rA, rC, rG or rU is found in a dinucleotide pair with any of the four dNMPs at neighboring upstream or downstream positions, and analyzed such preference. When the upstream dNMP is taken (NR), clear preferences exist around early-firing ARS’s in the *rnh201*-null libraries that are also specific on the leading or lagging strands. In particular, dArA, and in part dArC and dArG are stronger on the leading strand than the lagging strand for all *rnh201*-null libraries of wild-type polymerases prepared with all the three techniques (**Figure 5A**, dArA: P=1.82×10^−4^, dArC: P=1.07×10^−4^, dArG: P=3.46×10^−4^). On the contrary, dCrA, dCrC, dCrG are stronger on the lagging strand (**Figure 5A**, dCrA: P=2.58×10^−2^, dCrC: P=7.59×10^−3^, dCrG: P=4.66×10^−3^). We also examined whether there was any difference in the dinucleotide preference of incorporated rNMPs on the leading and lagging strands for the mutant DNA polymerase libraries. In the *pol2-M644G* mutant libraries, the preferred patterns are dArA, dArC, dArG, and dArU. These results show that in Pol ε mutant, the rNMPs tend to be incorporated after a dAMP, and this pattern is significantly stronger on the leading than on the lagging strand both for emRiboSeq and RHII-HydEn-seq libraries (**Figure 5B**, dArA: P=2.43×10^−4^, dArC: P=1.74×10^−4^, dArG: P=1.74×10^−4^, dArU: P=7.59×10^−3^). Differently, in the Pol δ mutant libraries (*pol3-L612M* emRiboSeq and *pol3-L612G* RHII-HydEn-seq libraries), the rNMPs tend to be incorporated after a dCMP, and this pattern is significantly stronger on the lagging strand (**Figure 5B**, dCrA: P=1.34×10^−3^, dCrC: P=5.51×10^−2^, dCrG: P=9.04×10^−3^, dCrU: P=8.99×10^−2^). Such different pattern is clearly evident by comparing side by side the heatmap data for the Pol ε and Pol δ mutants both for emRiboSeq and RHII-HydEn-seq libraries (**Figure 5B**). Hence, the pattern for the wild-type DNA polymerase libraries discussed above appears like a combination of the two patterns observed in Pol ε and Pol δ mutant libraries. The rNMPs after a dAMP are preferred on the leading strand, and the rNMPs after a dCMP are preferred on the lagging strand (**Figure 5A, B)**. Such leading and lagging patterns in wild-type polymerases and Pol ε and Pol δ mutants are clearly distinct in the early ARS’s, while for late ARS’s, the difference is maintained, although it is less evident (**Suppl Figure 5A, B**).

**Figure 5.**
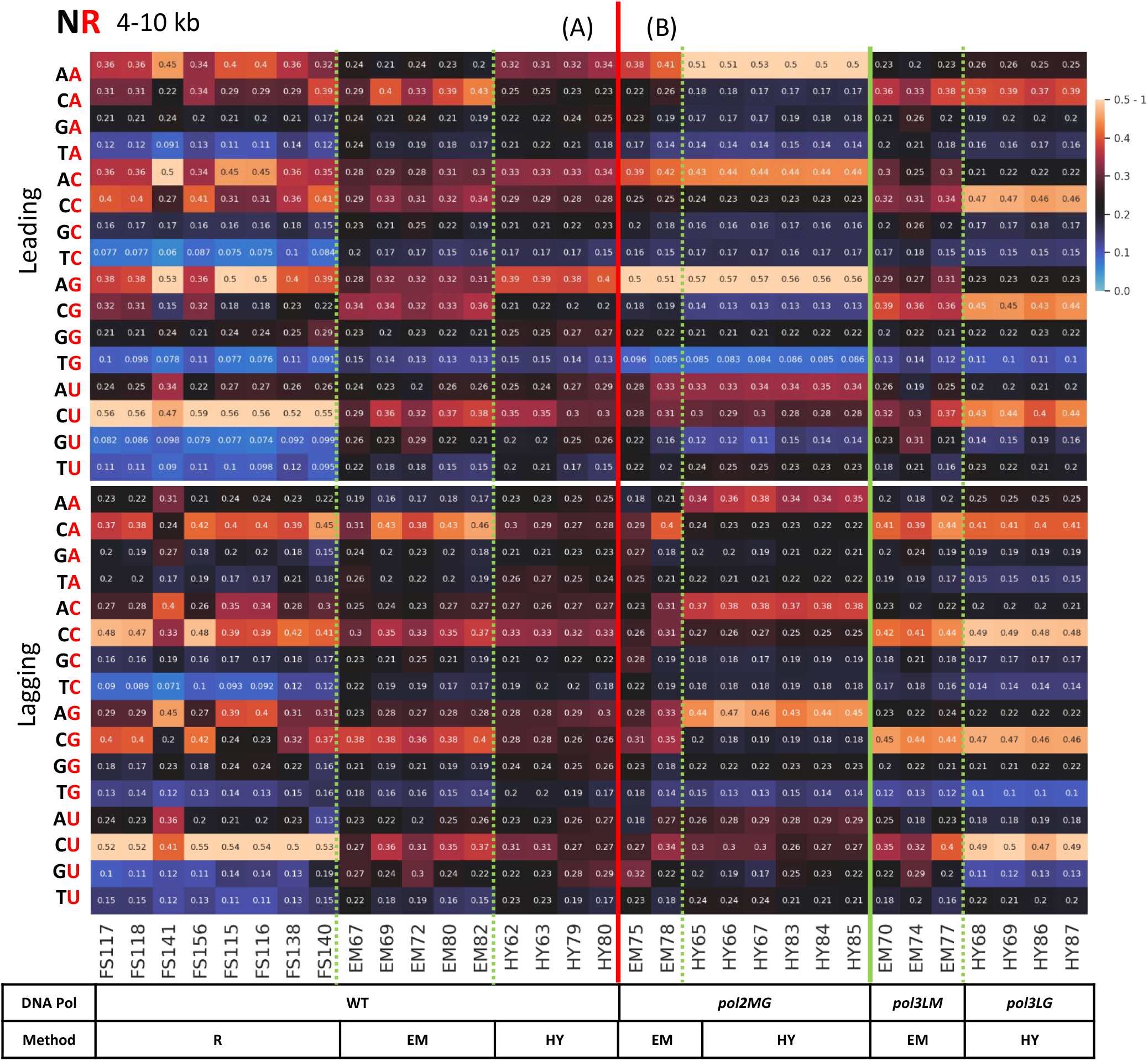
Marked dinucleotide NR preferences are revealed on the leading and lagging strands for wild-type and mutant Pols around 4-10 kb of early-firing ARS’s in *rnh201-*null libraries. Heatmap analyses with the normalized frequency of dinucleotides composed of the incorporated rNMP (R: rA, rC, rG, or rU) and its upstream neighbor (N: dA, dC, dG or dT) (NR) around early-firing ARS’s in *rnh201-*null libraries. The counts for each type of dinucleotide are normalized to the dinucleotide frequencies of the 4-10-kb window for the leading (top) or lagging (bottom) strand around early-firing ARS’s in the sacCer2 reference genome for all the ribose-seq and emRiboSeq libraries, and in the L03 reference genome for all the RHII-HydEn-seq libraries. The normalized frequency means the probability of an rNMP to be incorporated in the second position in the dinucleotide. The sum of four normalized frequencies with the same type of incorporated rNMP is further normalized to 1. Hence, 0.25 is the expected normalized frequency if there is no rNMP incorporation preference. The corresponding formula used is shown in the Methods section. The background nucleotide frequencies of ribose-seq and emRiboSeq (according to the sacCer2 reference genome), and RHII-HydEn-seq (according to the L03 reference genome) libraries are reported in **Supplementary Table 2A** and **2B**, respectively. The rNMP-incorporation position in the dinucleotide is shown in red at the top left of the heatmap. Each column of the heatmap shows the results of a specific library. The table underneath the heatmap shows the genotypes of DNA polymerase and the technique used for the rNMP library preparation. The thick, vertical, red line separates **(A)** data obtained from wild-type DNA polymerases from **(B)** data obtained with mutant DNA polymerases. The thick, vertical, green lines separate data obtained from wild-type DNA polymerases and from different mutant DNA polymerases libraries. The dashed, green lines separate data obtained using different rNMP mapping techniques. Each row shows results obtained for a type of rNMP. The bar to the right shows how normalized frequencies are represented as different colors: black for 0.25; black to yellow for 0.25 to 0.5 – 1, and black to light blue for 0.25 to 0. R, ribose-seq libraries; EM, emRiboSeq libraries; HY, RHII-HydEn-seq libraries; *pol2MG, pol2-M644G* mutant; *pol3LM,pol3-L612M* mutant for emRiboSeq libraries; *pol3LG, pol3-L612G* mutant for RHII-HydEn-seq libraries.

We performed a similar analysis for dinucleotides (NR) within the 0-200-nt window of the leading and lagging strands. The preferences of rNMP after a dAMP in the leading strand, and rNMP after a dCMP in the lagging strand also exist around early-firing ARS’s in the 0-200-nt window (**Figure 6A, B**). However, compared to the 4-10-kb window, the preference is weaker, and the bias of those patterns between the leading strand and lagging strand is also reduced. It is because the 0-200 nt corresponds to the DNA Pol δ to Pol ε-handoff phase, resulting in a combined Pol δ and Pol ε activity on the leading strand. To make a better comparison between the two different window lengths, we extracted the normalized frequencies of all 16 dinucleotides (NR) and drew box plots of preferred rNMP incorporation patterns on the leading and lagging strands in wild-type polymerase, and Pol ε and Pol δ mutant libraries. We find that in wild-type DNA polymerase libraries, all the dArN patterns are significantly preferred on the leading strand, and all dCrN patterns except dCrU are significantly preferred on the lagging strand within 4-10-kb window around early-firing ARS’s (**Figure 7A**). However, within the 0-200-nt window, dArC is the only pattern that is significantly preferred on the leading strand, and there is no significantly preferred pattern on the lagging strand (**Figure 7D**). In Pol ε mutant libraries, there are stronger preferences for all dArN patterns, and all preferences are significant within both 4-10 kb and 0-200-nt window. However, the *P* values are smaller within the 4-10-kb window (**Figure 7B, E**). In Pol δ mutant libraries, there are significant dCrA and dCrG preferences in the 4-10-kb window, and no significant preference is present within the 0-200-nt window (**Figure 7C, F**). These comparisons show that the preferred patterns are more significant within the 4-10-kb window, in which most of the Pol δ to ε switches already happened, compared to the 0-200-nt window. We also performed the same analysis in the 0-500 nt and 0-100-nt windows (**Supplementary Figure 8)**. Comparing the four windows, we find that the most significant preferences occur in the 4-10-kb window, then 0-500-nt window, then 0-200-nt window, and the 0-100-nt window has the least significant preference of patterns. This phenomenon suggests that there is less leading/lagging strand difference in the windows closer to the ARS, in which less Pol δ to Pol ε switch happened, and that there is more Pol δ activity on the leading strand in these windows. The same preferred rNMP patterns are also revealed, although less prominently, around the late-firing ARS’s, and the difference is more significant in the 4-10-kb window (**Supplementary Figure 7**,**9**).

**Figure 6.**
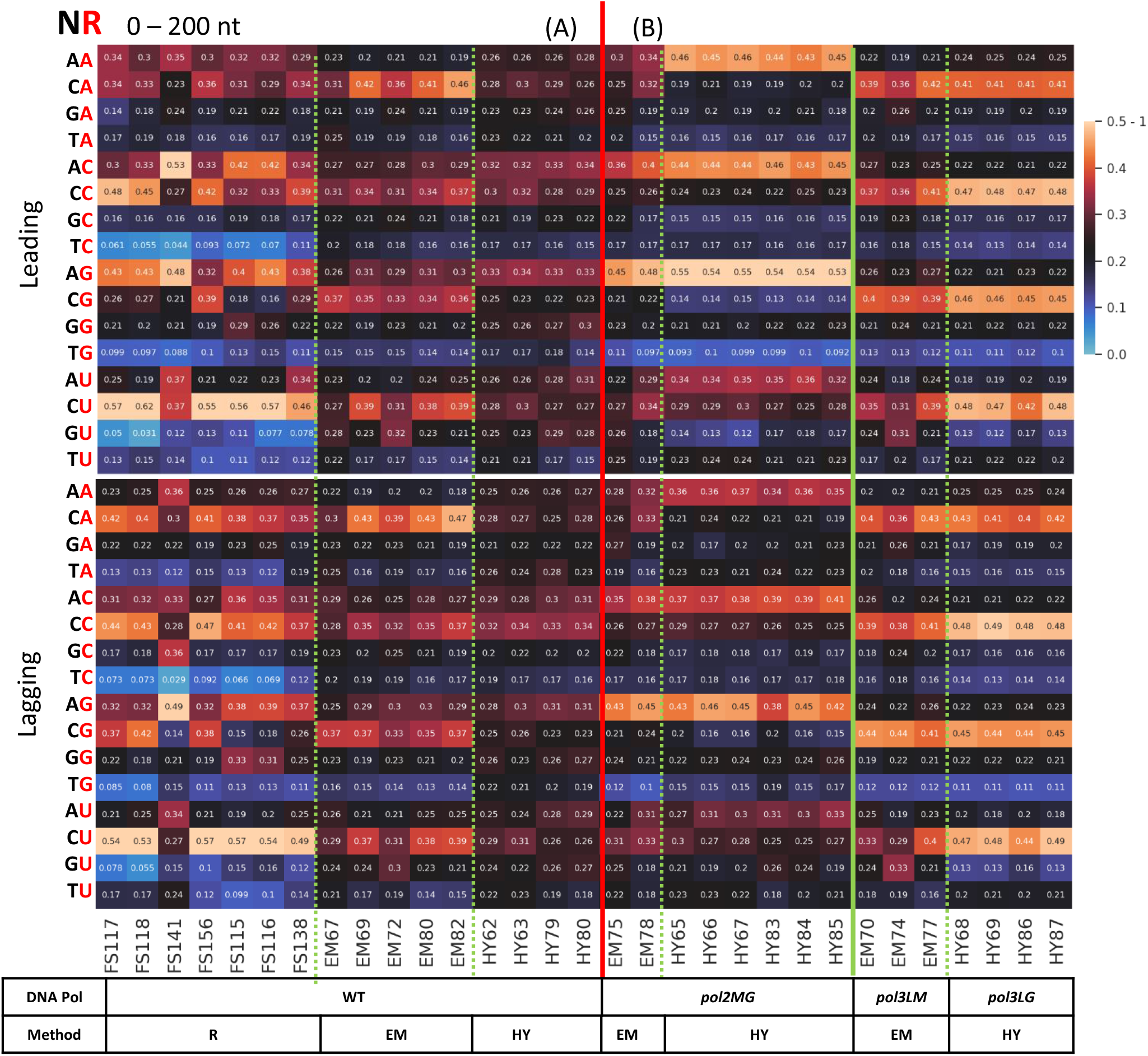
Dinucleotide (NR) preference of incorporated rNMPs on the leading and lagging strands is weakened around 0-200 nt of early-firing ARS’s in *rnh201-*null libraries. Heatmap analyses with the normalized frequency of dinucleotides composed of the incorporated rNMP (R: rA, rC, rG, or rU) and its upstream neighbor (N: dA, dC, dG or dT) (NR) around early-firing ARS’s in *rnh201-*null libraries. The counts for each type of dinucleotide are normalized to the dinucleotide frequencies of the 0-200-nt window for the leading (top) or lagging (bottom) strand around early-firing ARS’s in the sacCer2 reference genome for all the ribose-seq and emRiboSeq libraries, and in the L03 reference genome for all the RHII-HydEn-seq libraries. The normalized frequency means the probability of an rNMP to be incorporated in the second position in the dinucleotide. The sum of four normalized frequencies with the same type of incorporated rNMP is further normalized to 1. Hence, 0.25 is the expected normalized frequency if there is no rNMP incorporation preference. The corresponding formula used is shown in the Methods section. The background nucleotide frequencies of ribose-seq and emRiboSeq (according to the sacCer2 reference genome), and RHII-HydEn-seq (according to the L03 reference genome) libraries are reported in **Supplementary Table 2A** and **2B**, respectively. The rNMP-incorporation position in the dinucleotide is shown in red at the top left of the heatmap. Each column of the heatmap shows the results of a specific library. The table underneath the heatmap shows the genotypes of DNA polymerase and the technique used for the rNMP library preparation. The thick, vertical, red line separates **(A)** data obtained from wild-type DNA polymerases from **(B)** data obtained with mutant DNA polymerases. The thick, vertical, green lines separate data obtained from wild-type DNA polymerases and from different mutant DNA polymerases libraries. The dashed, green lines separate data obtained using different rNMP mapping techniques. Each row shows results obtained for a type of rNMP. The bar to the right shows how normalized frequencies are represented as different colors: black for 0.25; black to yellow for 0.25 to 0.5 – 1, and black to light blue for 0.25 to 0. R, ribose-seq libraries; EM, emRiboSeq libraries; HY, RHII-HydEn-seq libraries; *pol2MG, pol2-M644G* mutant; *pol3LM,pol3-L612M* mutant for emRiboSeq libraries; *pol3LG, pol3-L612G* mutant for RHII-HydEn-seq libraries.

**Figure 7.**
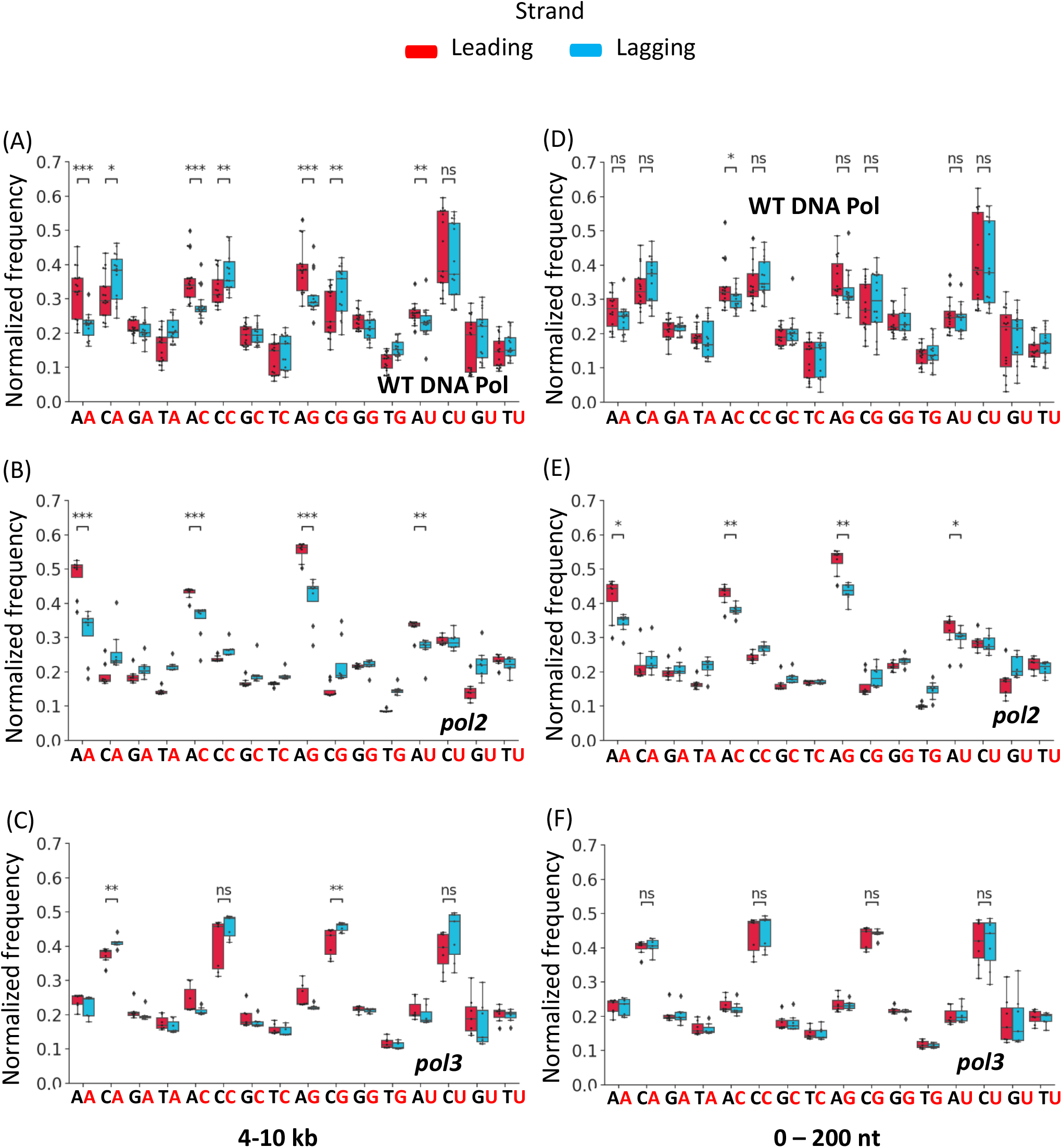
The rNMPs are preferentially preceded by dA on the leading and dC on the lagging strand within the 4-10 kb region around early-firing ARS’s in *rnh201*-null libraries. Boxplot of dinucleotide (NR) normalized frequency of each rNMP library around early-firing ARS’s. Outliers of 1.5 interquartile range (IQR) are marked as diamonds. The rNMP incorporation normalized frequencies in the 4-10-kb window of the leading or lagging strands around early-firing ARS are used in **A**-**C**, and the normalized frequencies of 0-200-nt window are used in **D**-**F**. Normalized dinucleotide (NR) frequencies of wild-type DNA polymerase libraries (N = 17) are shown in **A** and **D**, respectively. Normalized dinucleotide (NR) frequencies of *pol2* mutant libraries, including the *pol2-M644G* mutant of emRiboSeq and RHII-HydEn-seq libraries (N = 8) are shown in **B** and **E**, respectively. Normalized dinucleotide (NR) frequencies of *pol3* mutant libraries, including *pol3-L612M* mutant of emRiboSeq and *pol3-L612G* mutant of RHII-HydEn-seq libraries (N = 7) are shown in **C** and **F**, respectively. Mann-Whitney U tests are performed on dinucleotides (NR) with rA in *pol2* mutant libraries (**B** and **E**), rC in *pol3* mutant libraries (**C** and **F**), and rA or rC incorporated in wild-type DNA polymerase libraries (**A** and **D**). ns: P>0.05, *:0.05>P>0.01, **:0.01>P>0.001, ***:P<0.001.

If we combine the rNMPs with their downstream neighbor (RN) in the 4-10-kb window around early-firing ARS’s, no clear preference is found in the ribose-seq libraries. The rAdT in wild-type DNA Pols is the only preferred pattern in emRiboSeq libraries. The rNMPs are likely to be incorporated before dTMPs in RHII-HydEn-seq libraries, which may be caused by a technical factor in the construction of these libraries (**Supplementary Figure 10**). The RN patterns in the 0-200-nt window show high similarity to the patterns in the 4-10-kb window (**Supplementary Figure 11**). Around the late-firing ARS’s, rAdC is preferred in wild-type DNA polymerase and *pol2-M644G* RHII-HydEn-seq libraries, and the rAdA pattern is disfavored in wild-type DNA polymerase and *pol2-M644G* RHII-HydEn-seq libraries within the 4-10-kb window of the lagging strand (**Supplementary Figure 12**). These patterns do not exist in the 0-200-nt window around late-firing ARS’s, which show a pattern similar to early-firing ARS’s (**Supplementary Figure 13**).

In summary, while the composition of incorporated rNMPs is practically the same, apparent differences can be found in the dinucleotide preference of rNMPs on the leading and lagging strands. These differences are more significant in the mutant DNA polymerase libraries and around early-firing ARS’s. Moreover, these distinct patterns are more accentuated within the most distant (4-10-kb) window around ARS’s, in which more Pol δ to Pol ε handoff happened. Our results reveal specific features of rNMP incorporation by DNA Pol ε and Pol δ. Our findings also demonstrate that the composition of rNMPs present in yeast DNA is not dictated by the specific DNA polymerase enzymes that incorporate the rNMPs. However, the position of the rNMPs along the leading and lagging strands of yeast genomic DNA is markedly affected by the different DNA polymerase enzymes.

## DISCUSSION

### The function of DNA Pol δ on the leading strand synthesis and ARS annotation deviation

The function of Pol δ around the replication origin on the leading strand was previously observed from work in yeast (22, 31), and from *in vitro* work (21). By mapping rNMPs incorporated in nuclear DNA of budding and fission yeast using the RHII-HydEn-seq technique, Zhou et al., built a model supporting Pol δ function after Pol α in the starting zone during leading strand synthesis and identified 465 ARS’s in the L03 reference genome (12). Our study analyzed the incorporated rNMPs on the leading and lagging strand around ARS’s in 47 libraries of different genotypes and yeast strains prepared by three rNMP-mapping techniques. We characterized the Pol δ and Pol ε activities starting from the current known ARS’s in the sacCer2 and the L03 reference genomes. We also identified the changing phase of the leading/lagging ratio of rNMP incorporation in these libraries, represented by either increase or decrease of such ratio, which provides direct proof of a DNA Pol δ to Pol ε switch at the beginning of the leading strand synthesis in the budding yeast genome. The Pol δ to Pol ε handoff identified in our study is 4 kb in ribose-seq and emRiboSeq libraries and 1 kb in RHII-HydEn-seq libraries (**Figure 3**). At the beginning of the handoff range, Pol δ synthesizes the leading strand around the majority of ARS’s and Pol ε synthesizes the leading strand around the majority of ARS’s at the end of this range. Hence, the handoff range is longer than the Pol δ-synthesized tract length in Zhou et al. study (12), which matches the range of the extreme leading/lagging ratio that corresponds to the beginning of changing phase in our study. The gradient of increase or decrease is related to the frequencies of handoff events that happened at a given distance to the ARS’s. Abundant switches from Pol δ to Pol ε lead to a rapid increase or decrease of the leading/lagging ratio, which is around 2 kb for the ribose-seq and emRiboSeq libraries. The rapid changing phase of RHII-HydEn-seq libraries is within the first 500 nt from the ARS, pointing out that most of the Pol δ to Pol ε switches happen in the first 500 nt on the leading strand synthesis in these libraries (**Figure 3 RHII-HydEn-seq zoom-in**). In the ribose-seq and emRiboSeq libraries, the deviation from the ARS annotation leads to a larger smoothing of the leading/lagging ratio for the combined ARS’s and a longer changing phase. RHII-HydEn-seq data with the ARS annotation in the L03 reference genome are more accurate than ribose-seq and emRiboSeq data with OriDB annotation in sacCer2 reference genome. The same genome identity between the RHII-HydEn-seq libraries and the L03 reference genome likely contributes to the more accurate results. Interestingly, we find that the most rapid changing phase is longer around the late than the early-firing ARS’s in the wild-type DNA polymerase libraries but not in the Pol ε and Pol δ mutant libraries, suggesting that the length of the Pol δ-synthesized tract before the handoff to Pol ε can be affected by the firing time only in the wild-type DNA polymerase libraries (**Figure 3**).

### Different DNA polymerase usage leads to different dinucleotide preference pattern on the leading and lagging strands

We discovered that rNMP incorporation in DNA sequences around the ARS’s that are synthesized by different replicative DNA polymerases displays distinct sequence contexts, represented by different patterns on the leading and lagging strands. Our study demonstrates that the composition of rNMPs in *rnh201*-null cells is strikingly similar on the leading and lagging strand within a 10-kb flank around the ARS’s in libraries with different DNA Pol δ and Pol ε alleles (wild-type or mutants). And the composition is even conserved with the rNMP composition in the full genome for the corresponding libraries (24) (**Figure 4, Supplementary Figure 4**). However, the dinucleotide preference of rNMP markedly changes on the leading and lagging strands at different distances from the ARS’s and in different DNA polymerase alleles. The dinucleotide preference in the 4-10-kb window on the leading strand, in which most Pol δ to Pol ε switches happens, is different from the preference observed in the same 4-10-kb window on the lagging strand. In this window, the leading strand is mainly synthesized by Pol ε, and the lagging strand is mainly synthesized by Pol δ. We discovered that DNA Pol ε tends to incorporate rNMPs after a dAMP, while DNA Pol δ tends to incorporate rNMPs after a dCMP in the 4-10-kb window (**Figure 5, Supplementary Figure 5**). The frequency of the preferred dinucleotide pattern is also different on the leading and lagging strand due to the different DNA polymerase usage. The dArN pattern of Pol ε is stronger on the leading strand since Pol ε synthesizes the main part of the leading strand, and the dCrN pattern of Pol δ is stronger in the lagging strand since Pol δ synthesizes the main part of lagging strand. These dArN-leading and dCrN-lagging patterns are particularly accentuated in the Pol ε and Pol δ mutant libraries, respectively (**Figure 5B**). In the wild-type polymerase libraries, we instead found a combination of the two patterns observed in Pol ε and Pol δ mutant libraries. We further checked if the frequency of the preferred pattern was significantly different in the four different windows, 4-10 kb, 0-500 nt, 0-200nt, and 0-100 nt on the leading and lagging strands. The leading/lagging strand differences of the preferred patterns show a progressive reduction of the significance with the shortening distance to the ARS’s. These results suggest that the DNA polymerase usage on the leading and lagging strands is more similar in the windows that are closer to the ARS’s location (**Figure 7, Supplementary Figure 7**,**8**,**9**). The similar DNA polymerase usage in the windows close to the ARS’s reveals the Pol δ activity on the leading strand, which is reduced with the occurrence of the Pol δ to Pol ε switch.

### Conclusion

By analyzing the characteristics of rNMP incorporation from 47 libraries of 13 different *Saccharomyces cerevisiae* genotypes and 5 different yeast strains prepared using 3 different techniques: ribose-seq, emRiboSeq, and RHII-HydEn-seq, we discovered specific preferences of rNMP incorporation on the leading strand in wild-type DNA polymerase and *pol2-M644G* libraries, and on the lagging strand in *pol1* and *pol3* mutant libraries (*pol1-L868M* and *pol3-L612M* for emRiboSeq libraries, *pol1-Y869A* and *pol3-L612G* for RHII-HydEn-seq libraries). We found that such preferences are weakened with the increasing ARS firing time. We also developed a model for the rate change of rNMP incorporation during DNA replication in yeast, which validates the fact that Pol δ also contributes to the early phase synthesis of the leading strand. Furthermore, we found distinct dinucleotide preferences of rNMP incorporation on the leading and lagging strands, which are generated by the different DNA polymerase usage on the leading and lagging strands. The different leading/lagging patterns of rNMP distribution are less distinct at the beginning of DNA replication, suggesting a similar DNA polymerase usage on the leading and lagging strands, and thus validating the fact that Pol δ contributes to the early phase synthesis of the leading strand.

## DATA AVAILABILITY

The *ribose-seq*, emRiboSeq, and RHII-HydEn-seq libraries used in the study are download in the NCBI BioProject under the accession number PRJNA613920, PRJNA271170, and PRJNA517710.

The only *pol2* mutant library of *ribose-seq* is under the NCBI BioProject accession number PRJNA261234.

The liftover software, bigWigToBedGraph software and the chain file from sacCer1 to sacCer2 are downloaded from UCSC Genome Browser Utilities (http://hgdownload.soe.ucsc.edu/downloads.html)

Ribose-Map software is stored in Github (https://github.com/agombolay/ribose-map). RibosePreferenceAnalysis tools deposited in Github are also used to generate heatmaps (https://github.com/xph9876/RibosePreferenceAnalysis). All other scripts are deposit in the Github (https://github.com/xph9876/rNMP_ARS_analysis).

## ACKNOWLEDGEMENT

We thank A. Gombolay for computational support, the Partnership for an Advanced Computing Environment (PACE) at the Georgia Institute of Technology for their research cyberinfrastructure resources and services, and all members of the Storici laboratory for assistance and feedback on this study. We acknowledge funding from the National Institutes of Health R01 ES026243 (F.S.) and the Howard Hughes Medical Institute (Faculty Scholar grant 55108574 to F.S.) for supporting this work.

