## Supplementary Figures 1-13 for "Ribonucleotide incorporation characteristics around yeast autonomously replicating sequences reveal the labor division of replicative DNA polymerases"

**Supplementary Table 1. All rNMP libraries used in the study.** All the libraries used in the study with their SRR Accession and BioProject Accession are listed. For the ribose-seq and emRiboSeq libraries, FASTQ files are downloaded, and Ribose-Map software is used to locate the rNMP incorporation in SGD sacCer2 reference genome. For the RHII-HydEn-seq libraries, aligned BigWig files with the L03 reference genome are used and converted into BED format.

**Supplementary Table 2. Background frequency of heatmaps in the *S. cerevisiae* sacCer2 and L03 reference genome.** Count and frequencies (%) of mononucleotides and dinucleotides of the 4-10 kb and 0-200-nt windows of the leading and lagging strands around early and late-firing ARS's in **(A)** sacCer2 and **(B)** L03 reference genome. The raw number of each dNMP or dinucleotide in the reference genome is counted, and the percentage is calculated. For the dinucleotide tables, "R" means incorporated ribonucleotide, "N" means deoxyribonucleotide neighbor of base A, C, G or T. The percentage is calculated by dividing the raw number of NR or RN to the sum of the four combinations for each fixed R., E.g., in the NR = AC, the percentage is ACx100/(AC+CC+GC+TC); for RN = AC, the percentage is ACx100/(AA+AC+AG+AT).

**Supplementary Figure 1. Model of rNMP incorporation around the ARS region.** Model of labor division by yeast replicative DNA polymerases and rNMP incorporation at the replication fork. DNA replication starts from the ARS, and the leading and lagging strands are synthesized at the ARS flanks. The leading strand synthesis starts upstream of the ARS center with a "lagging strand" primer, which is initiated by DNA Pol α and extended by Pol δ. Pol ε takes over and synthesize the bulk of the leading strand afterward. Finally, DNA Pol ε hands back to Pol δ when the leading strand collides with the incoming lagging strand (12, 21). The lagging strand is composed of Okazaki fragments, which are synthesized by Pol α and Pol δ. Abundant rNMPs (R, in purple) are misincorporated into the newly synthesized strands during the replication process.

**Supplementary Figure 2. The rNMP incorporation rate on the leading and lagging strands.** The rNMP counts are divided by background dNMP frequency to obtain the rNMP per base (RPB) value, which is the direct measurement of the rNMP-incorporation rate. The average RPB values in each 0.5 kb bin (0.1 kb for zoom-in) for the first 15 kb of the leading and lagging strands (1 kb for zoom-in) are calculated with maximum likelihood estimation and shown in the plots. R, ribose-seq libraries; EM, emRiboSeq libraries; HY, RHII-HydEn-seq libraries; *rnh201*, RNase H2 defective mutant; *pol2, pol2-*

*M644G* mutant; *pol3, pol3-L612G* mutant for RHII-HydEn-seq libraries and *pol3-L612M* mutant for emRiboSeq libraries. **(A)** The RPB value changes on the leading strands. **(B)** The RPB value changes on the leading strands.

**Supplementary Figure 7. The dinucleotide (NR) preference of incorporated rNMPs is different on the leading and lagging strand around late-firing ARS's.** Boxplot of dinucleotide (NR) normalized frequency of each rNMP library around late-firing ARS's. Outliers of 1.5 interquartile range (IQR) are marked as diamonds. The normalized frequencies of rNMP incorporation in the 4-10-kb window of the leading or lagging strand around late-firing ARS are used in **A**-**C**, and the normalized frequencies in the 0-200-nt window are used in **D**-**F**. Normalized dinucleotide (NR) frequency of wild-type DNA polymerase libraries (N = 17) is shown in **A** and **D**, respectively. Normalized dinucleotide (NR) frequency of *pol2* mutant libraries, including the *pol2-M644G* mutant of emRiboSeq and RHII-HydEn-seq libraries (N = 8) are shown in **B** and **E**, respectively. Normalized dinucleotide (NR) frequency of *pol3* mutant libraries, including *pol3-L612M* mutant of emRiboSeq and *pol3-L612G* mutant of RHII-HydEn-seq libraries (N = 7) are shown in **C** and **F**, respectively. Mann-Whitney U tests are performed on dinucleotides (NR) with rA in *pol2* mutant libraries (**B** and **E**), rC in *pol3* mutant libraries **(C** and **F**), and rA or rC incorporated in wild-type DNA polymerase libraries (**A** and **D**). ns: P>0.05, *:0.05>P>0.01, **:0.01>P>0.001, ***:P<0.001.

**Supplementary Figure 8. The dinucleotide (NR) preference of incorporated rNMPs within the 0-500 nt and 0-100-nt windows around early-firing ARS's.** Boxplot of dinucleotide (NR) normalized frequency of each rNMP library around early-firing ARS's. Outliers of 1.5 interquartile range (IQR) are marked as diamonds. The normalized frequencies of rNMP incorporation in the 0-500-nt window of the leading or lagging strand around early-firing ARS are used in **A**-**C**, and the normalized frequencies in the 0-100-nt window are used in **D**-**F**. Normalized dinucleotide (NR) frequency of wild-type DNA polymerase libraries (N = 16) is shown in **A** and **D**, respectively. Normalized dinucleotide (NR) frequency of *pol2* mutant libraries, including the *pol2-M644G* mutant of emRiboSeq and RHII-HydEn-seq libraries (N = 8) are shown in **B** and **E**, respectively. Normalized dinucleotide (NR) frequency of *pol3* mutant libraries, including *pol3-L612M* mutant of emRiboSeq and *pol3-L612G* mutant of RHII-HydEn-seq libraries (N = 7) are shown in **C** and **F**, respectively. Mann-Whitney U tests are performed on dinucleotides (NR) with rA in *pol2* mutant libraries (**B** and **E**), rC in *pol3* mutant libraries **(C** and **F**), and rA or rC incorporated in wild-type DNA polymerase libraries (**A** and **D**). ns: P>0.05, *:0.05>P>0.01, **:0.01>P>0.001, ***:P<0.001.

**Supplementary Figure 9. The dinucleotide (NR) preference of incorporated rNMPs within the 0-500 nt and 0-100-nt windows around late-firing ARS's.** Boxplot of dinucleotide (NR) normalized frequency of each rNMP library around late-firing ARS's. Outliers of 1.5 interquartile range (IQR) are marked as diamonds. The normalized frequencies of rNMP incorporation in the 0-500-nt window of the leading or lagging strand around late-firing ARS are used in **A**-**C**, and the normalized frequencies in the 0-100-nt window are used in **D**-**F**. Normalized dinucleotide (NR) frequency of wild-type DNA polymerase libraries (N = 16) is shown in **A** and **D**, respectively. Normalized dinucleotide (NR) frequency of *pol2* mutant libraries, including the *pol2-M644G* mutant of emRiboSeq and RHII-HydEn-seq libraries (N = 8) are shown in **B** and **E**, respectively. Normalized dinucleotide (NR) frequency of *pol3* mutant libraries, including *pol3-L612M* mutant of emRiboSeq and *pol3-L612G* mutant of RHII-HydEn-seq libraries (N = 7) are shown in **C** and **F**, respectively. Mann-Whitney U tests are performed on dinucleotides (NR) with rA in *pol2* mutant libraries (**B** and **E**), rC in *pol3* mutant libraries **(C** and **F**), and rA or rC incorporated in wild-type DNA polymerase libraries (**A** and **D**). ns: P>0.05, *:0.05>P>0.01, **:0.01>P>0.001, ***:P<0.001.

**Supplementary Figure 13 Dinucleotide (RN) preference of incorporated rNMPs within the 0-200-nt window around late-firing ARS's in *rnh201*-null libraries.** Heatmap analyses with the normalized frequency of dinucleotides composed of the incorporated rNMP (R: rA, rC, rG, or rU) and its downstream neighbor (N: dA, dC, dG or dT) (RN) around late-firing ARS's in *rnh201*-null libraries. The counts for each type of dinucleotide are normalized to the dinucleotide frequencies of the 0-200-nt window for the leading (top) or lagging (bottom) strand around late-firing ARS's in the sacCer2 reference genome for all the ribose-seq and emRiboSeq libraries, and in the L03 reference genome for all the RHII-HydEn-seq libraries. The normalized frequency means the probability of an rNMP to be incorporated in the first position in the dinucleotide. The sum of four normalized frequencies with the same type of incorporated rNMP is further normalized to 1. Hence, 0.25 is the expected normalized frequency if there is no rNMP incorporation preference. The corresponding formula used is shown in the Methods section. The background nucleotide frequencies of ribose-seq and emRiboSeq (according to the sacCer2 reference genome), and RHII-HydEn-seq (according to the L03 reference genome) libraries are reported in **Supplementary Table 2A** and **2B**, respectively. The rNMP incorporation position in the dinucleotide is shown in red at the top left of the heatmap. Each column of the heatmap shows the results of a specific library. The table underneath the heatmap shows the genotypes of DNA polymerase and the technique used for the rNMP library preparation. The thick, vertical, red line separates **(A)** data obtained from wild-type DNA polymerases from **(B)** data obtained with mutant DNA polymerases. The thick, vertical, green lines separate data obtained from wild-type DNA polymerases and from different mutant DNA polymerases libraries. The dashed, green lines separate data obtained using different rNMP mapping techniques. Each row shows results obtained for a type of rNMP. The bar to the right shows how normalized frequencies are represented as different colors: black for 0.25; black to yellow for 0.25 to 0.5 – 1, and black to light blue for 0.25 to 0. R, ribose-seq libraries; EM, emRiboSeq libraries; HY, RHII-HydEn-seq libraries; *pol2MG, pol2-M644G* mutant; *pol3LM,pol3-L612M* mutant for emRiboSeq libraries; *pol3LG, pol3-L612G* mutant for RHII-HydEn-seq libraries.

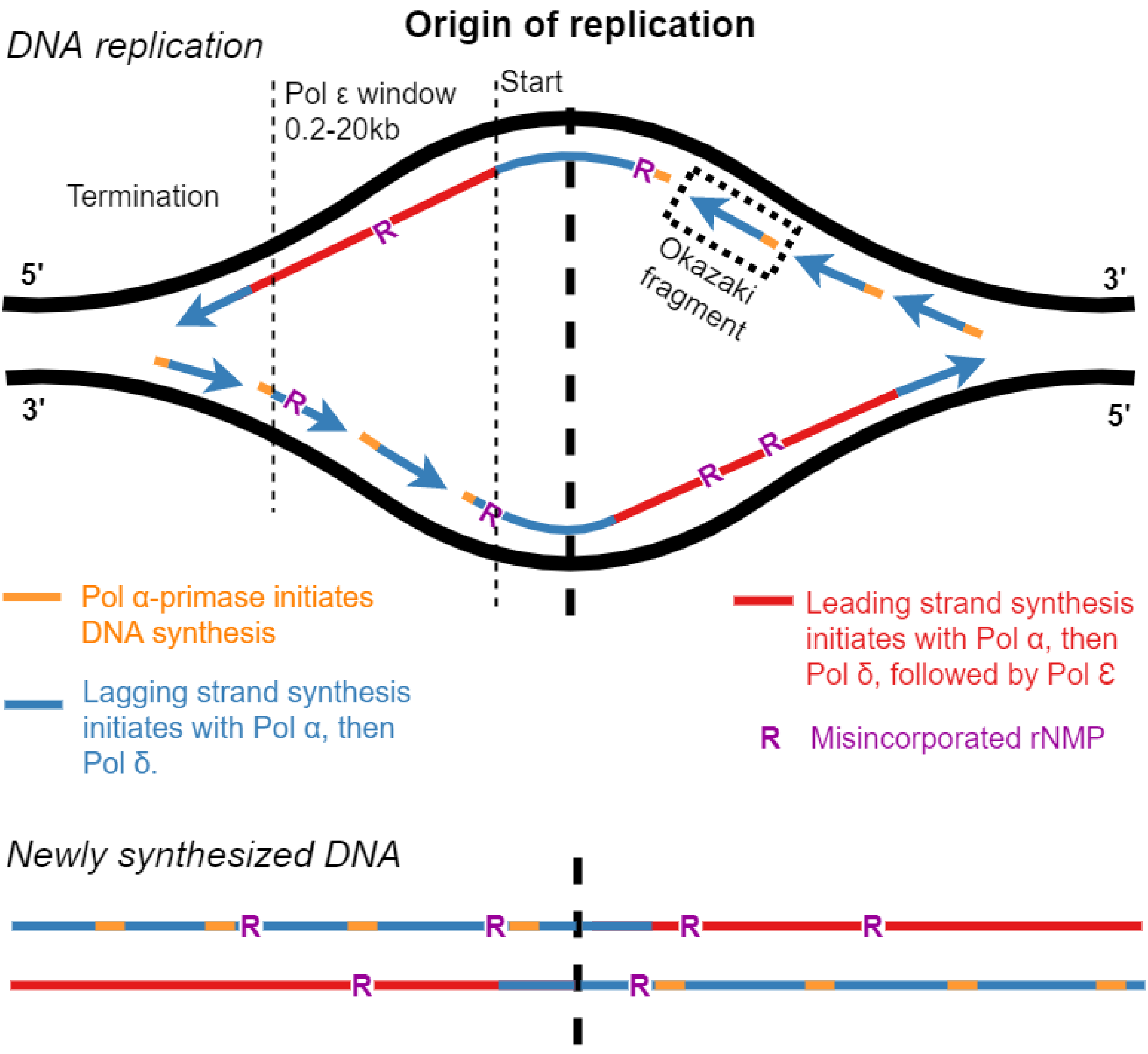


**Supplementary Figure 1**

**Supplementary Figure 2**

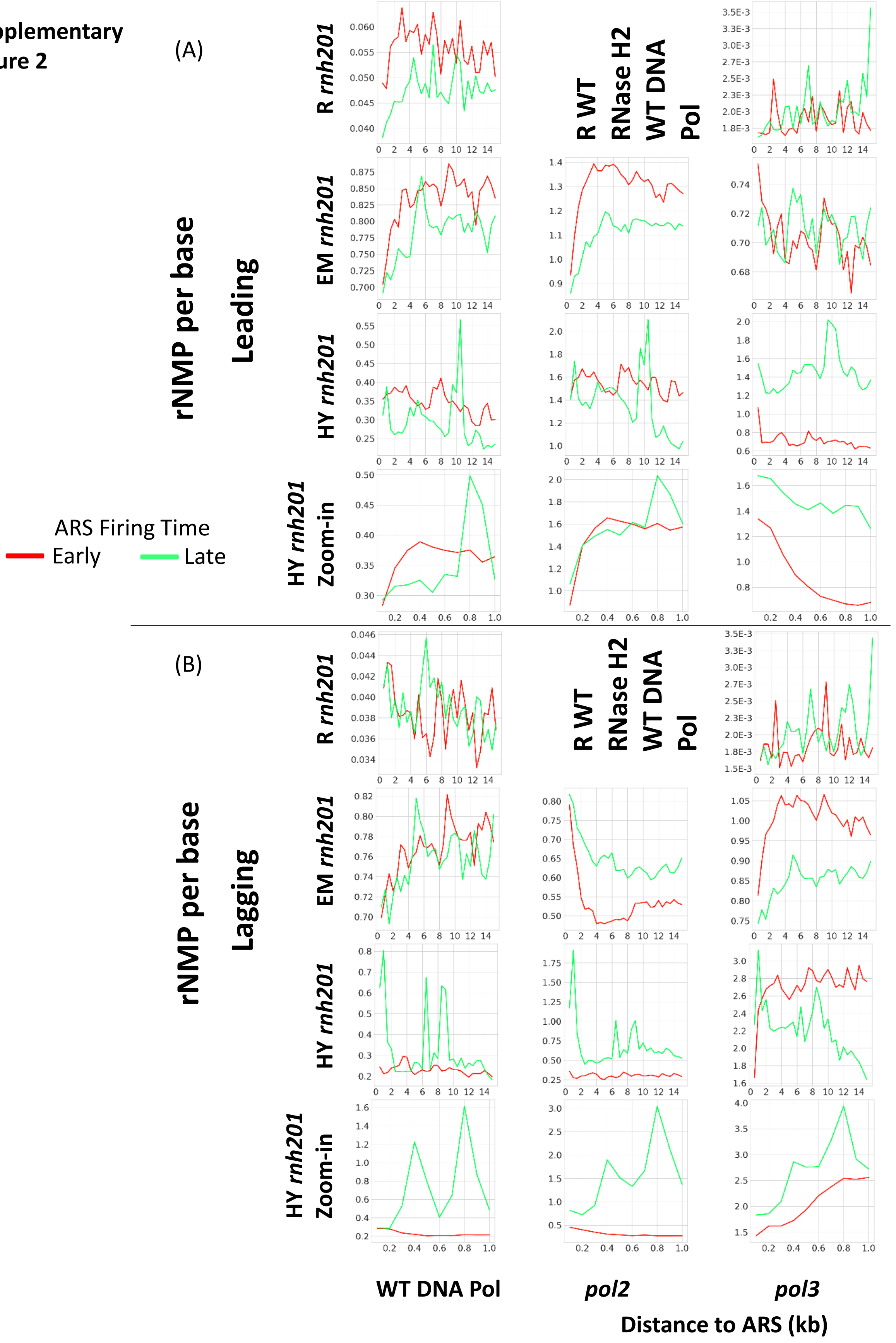

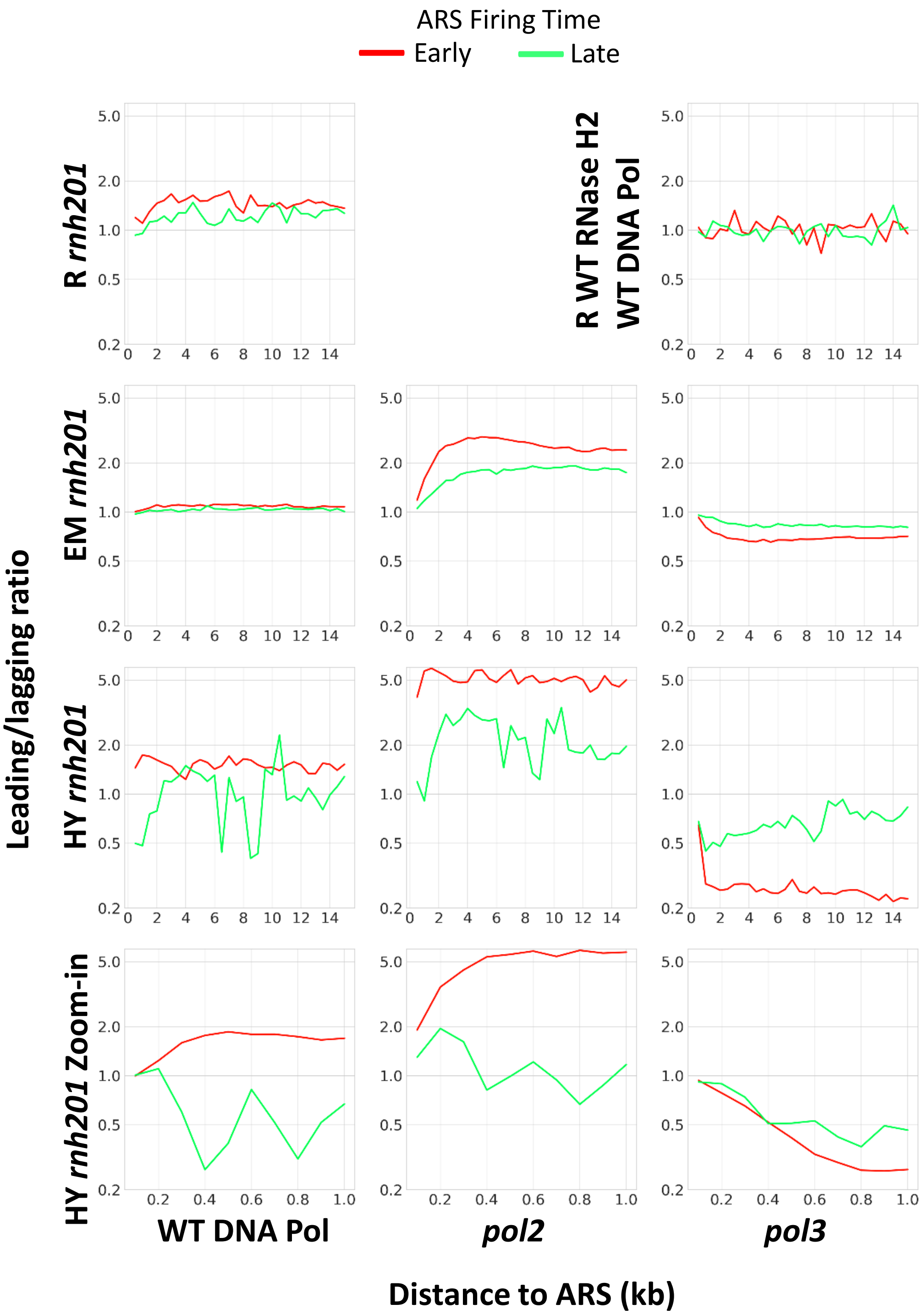


**Supplementary Figure 3**

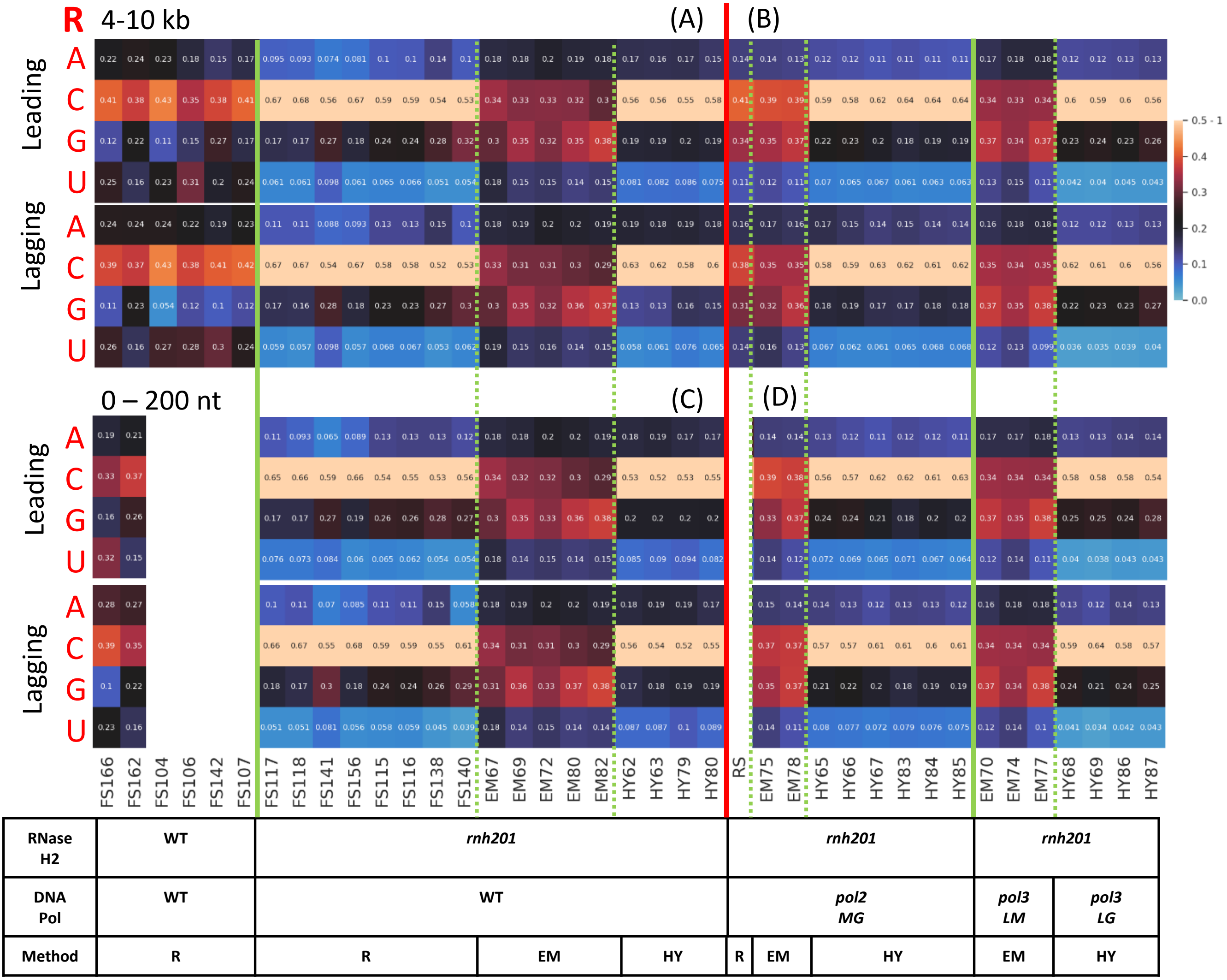


**Supplementary Figure 4**

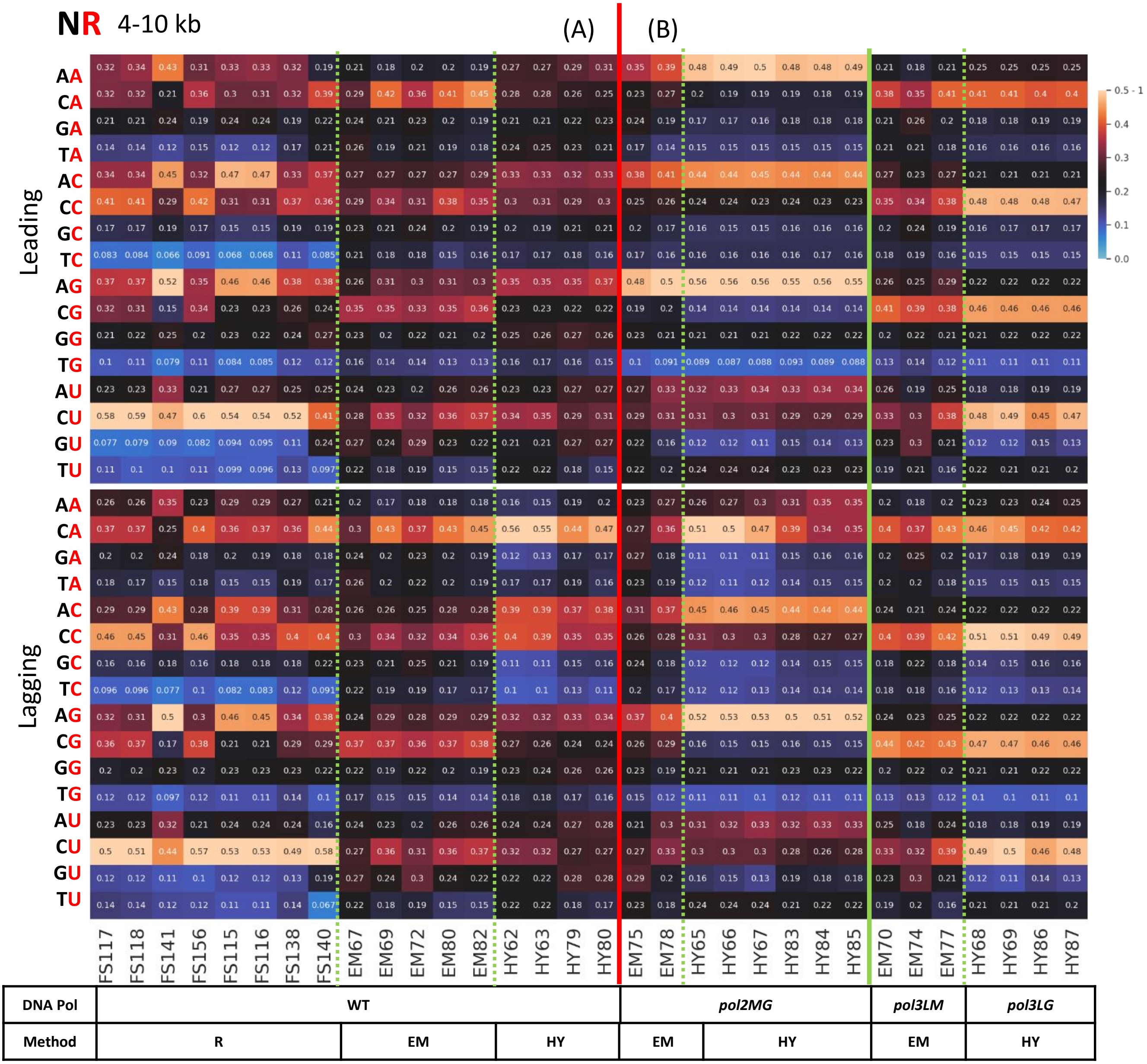


**Supplementary Figure 5**

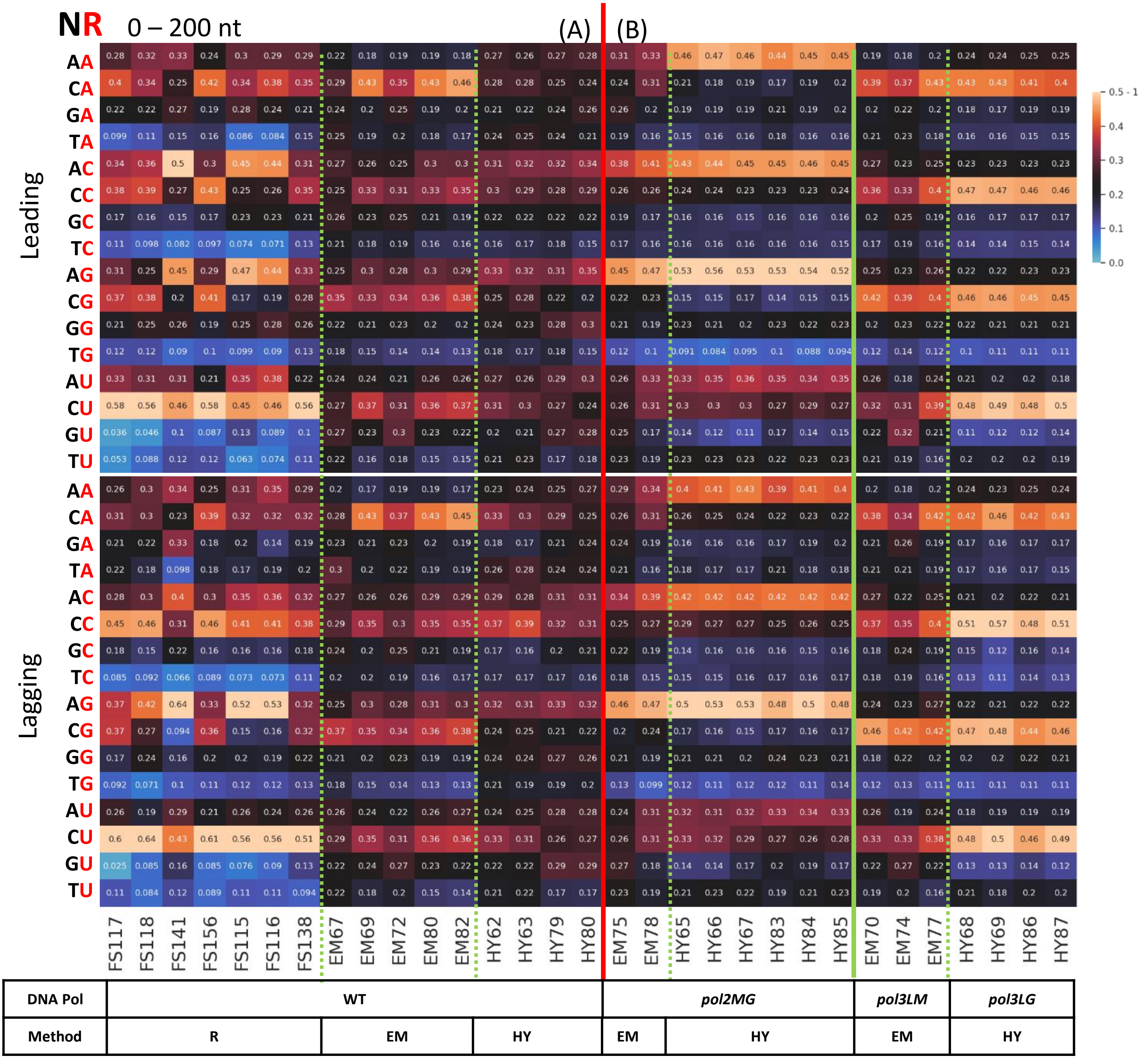


**Supplementary Figure 6**

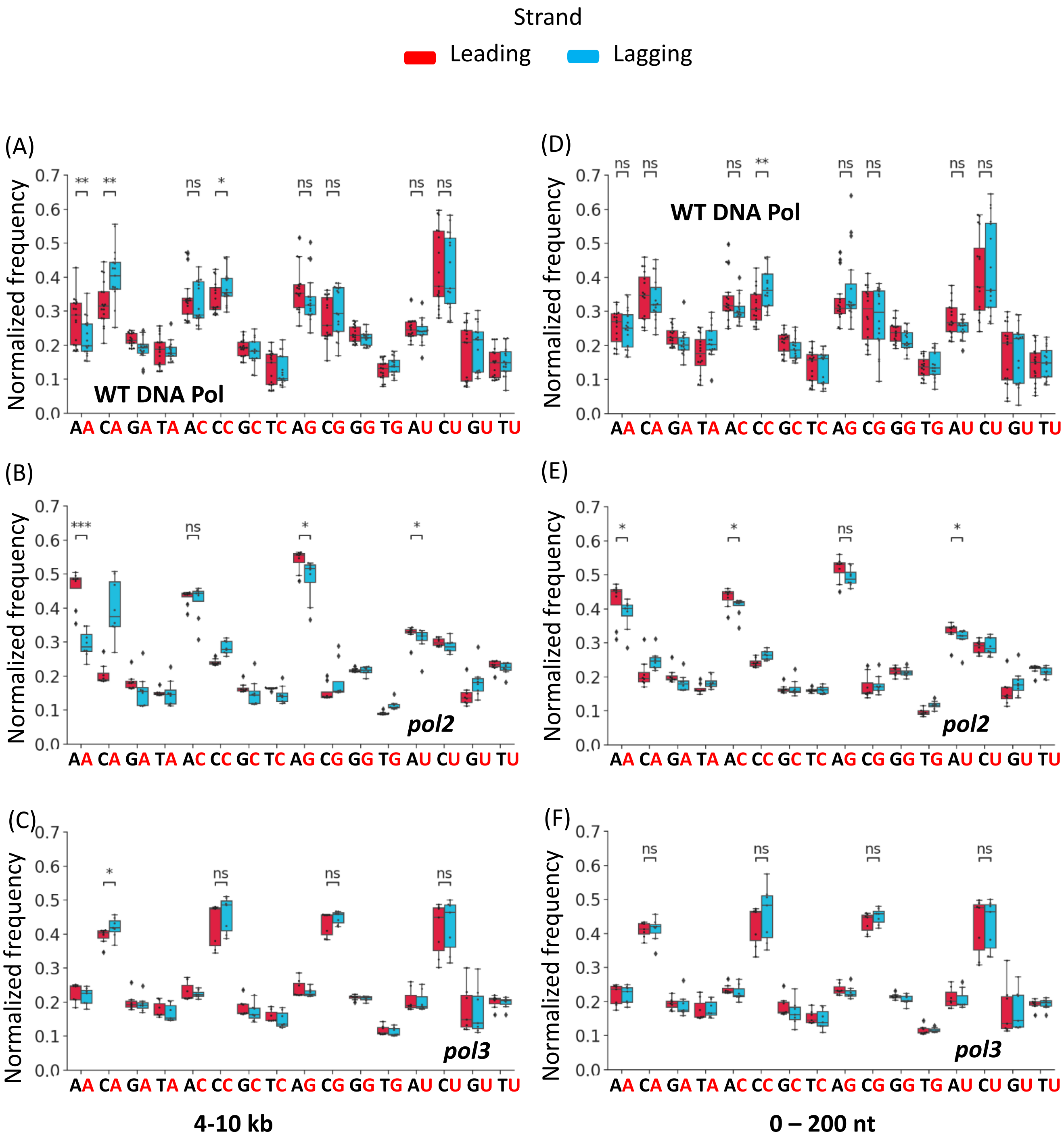


**Supplementary Figure 7**

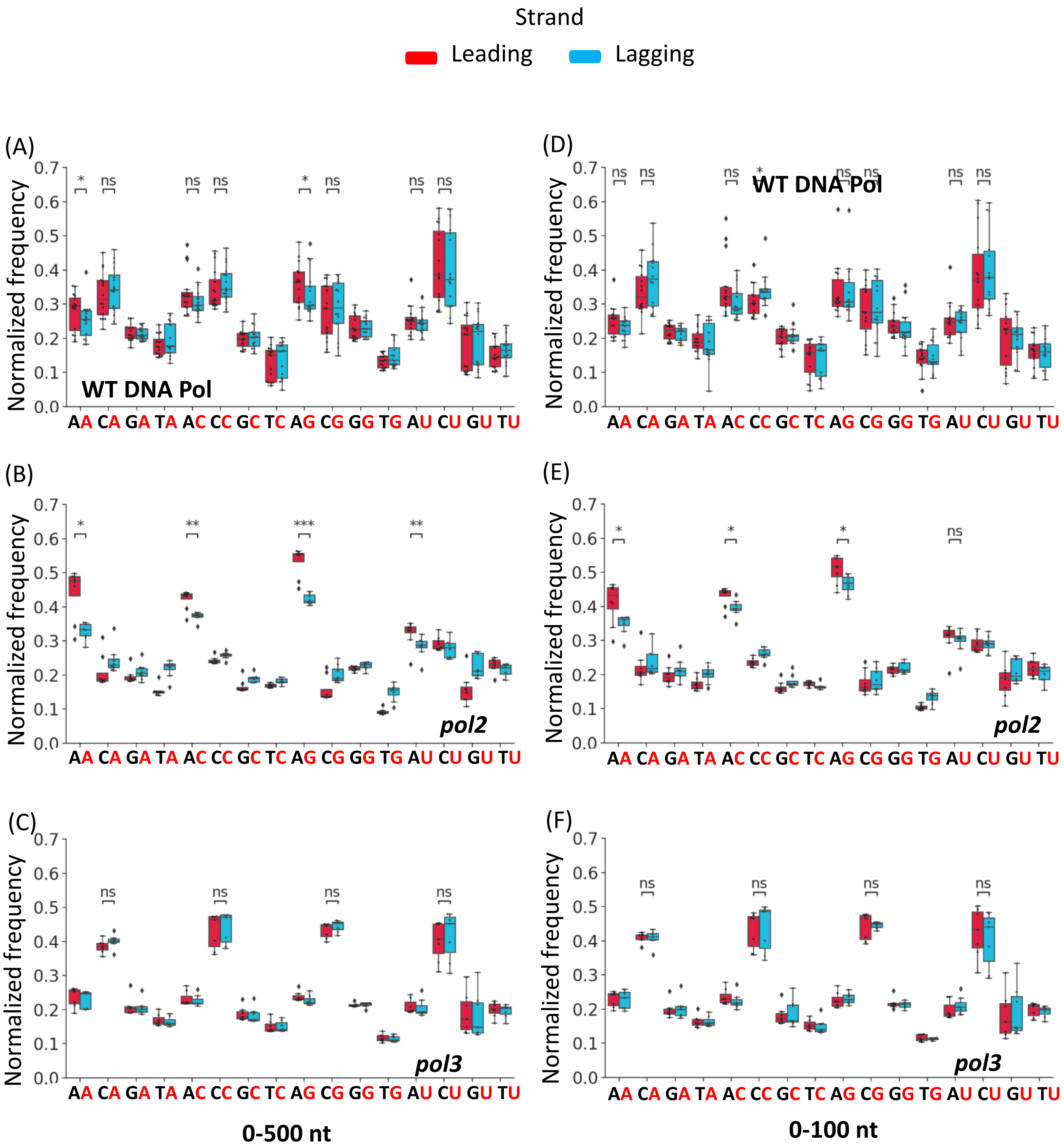


**Supplementary Figure 8**

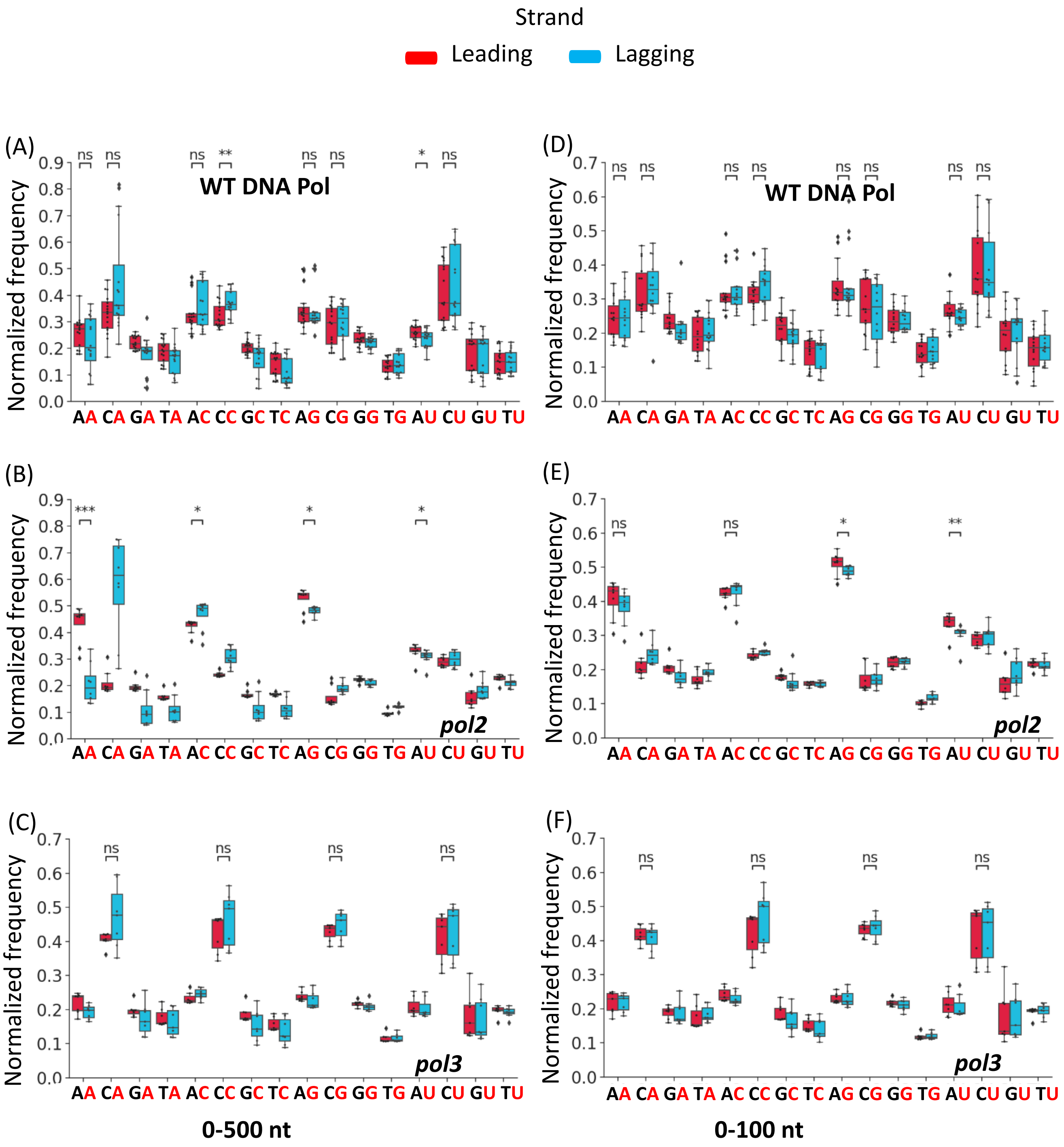


**Supplementary Figure 9**

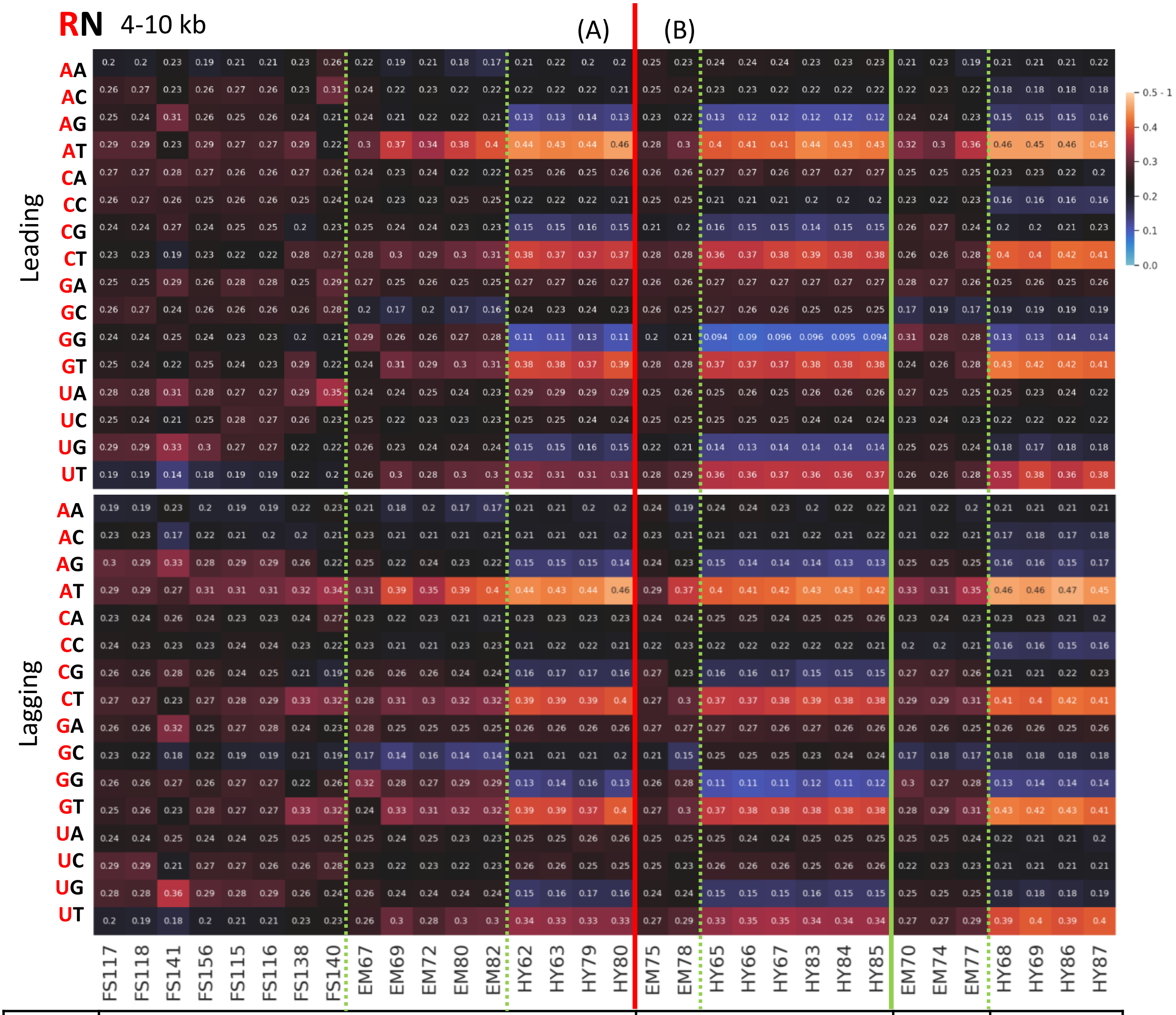


| DNA Pol | WT | | | *pol2MG* | | *pol3LM* | *pol3LG* |
|---|---|---|---|---|---|---|---|
| Method | R | EM | HY | EM | HY | EM | HY |

**Supplementary Figure 10**

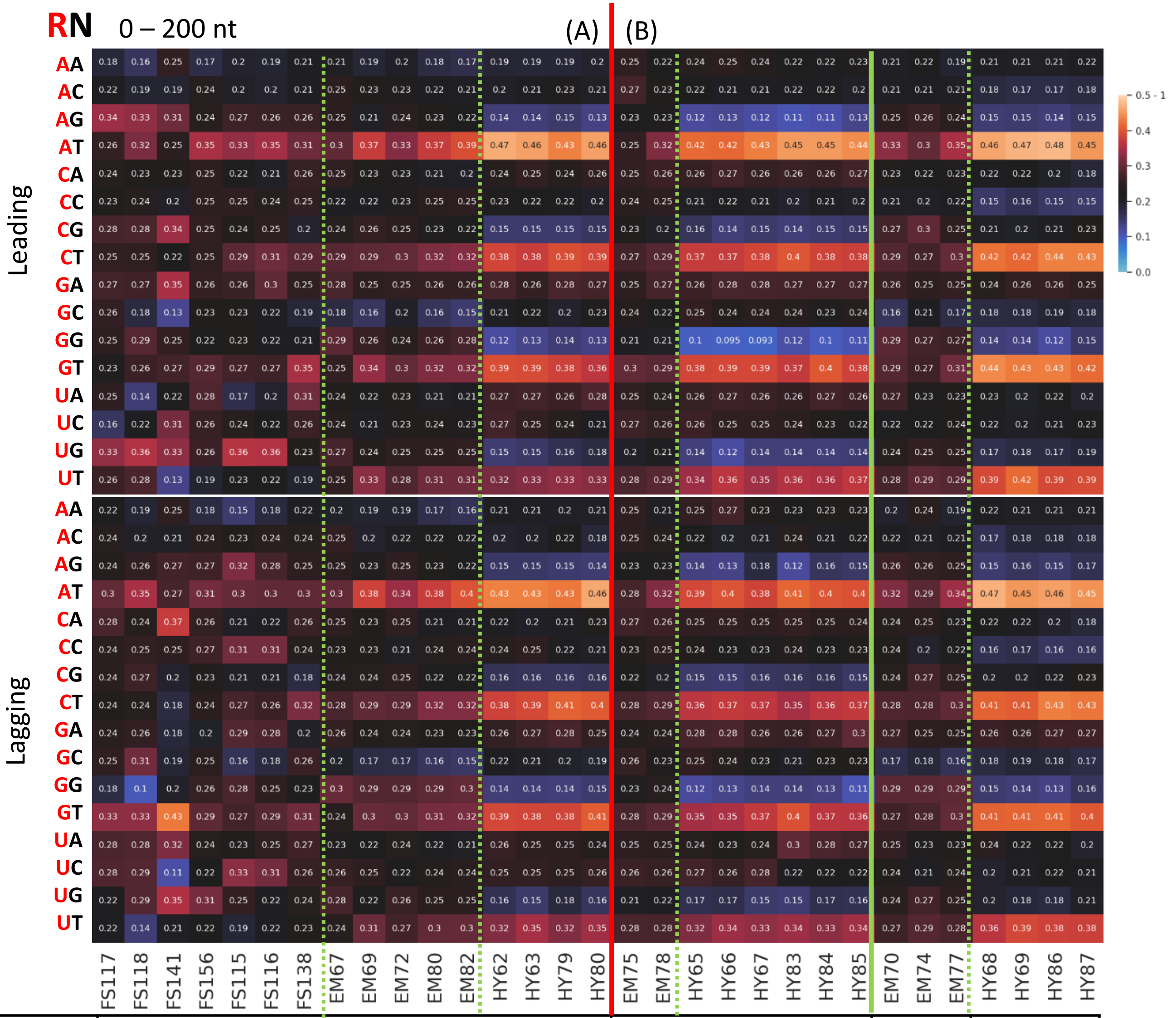


| DNA Pol | WT | | | *pol2MG* | | *pol3LM* | *pol3LG* |
|---|---|---|---|---|---|---|---|
| Method | R | EM | HY | EM | HY | EM | HY |

**Supplementary Figure 11**

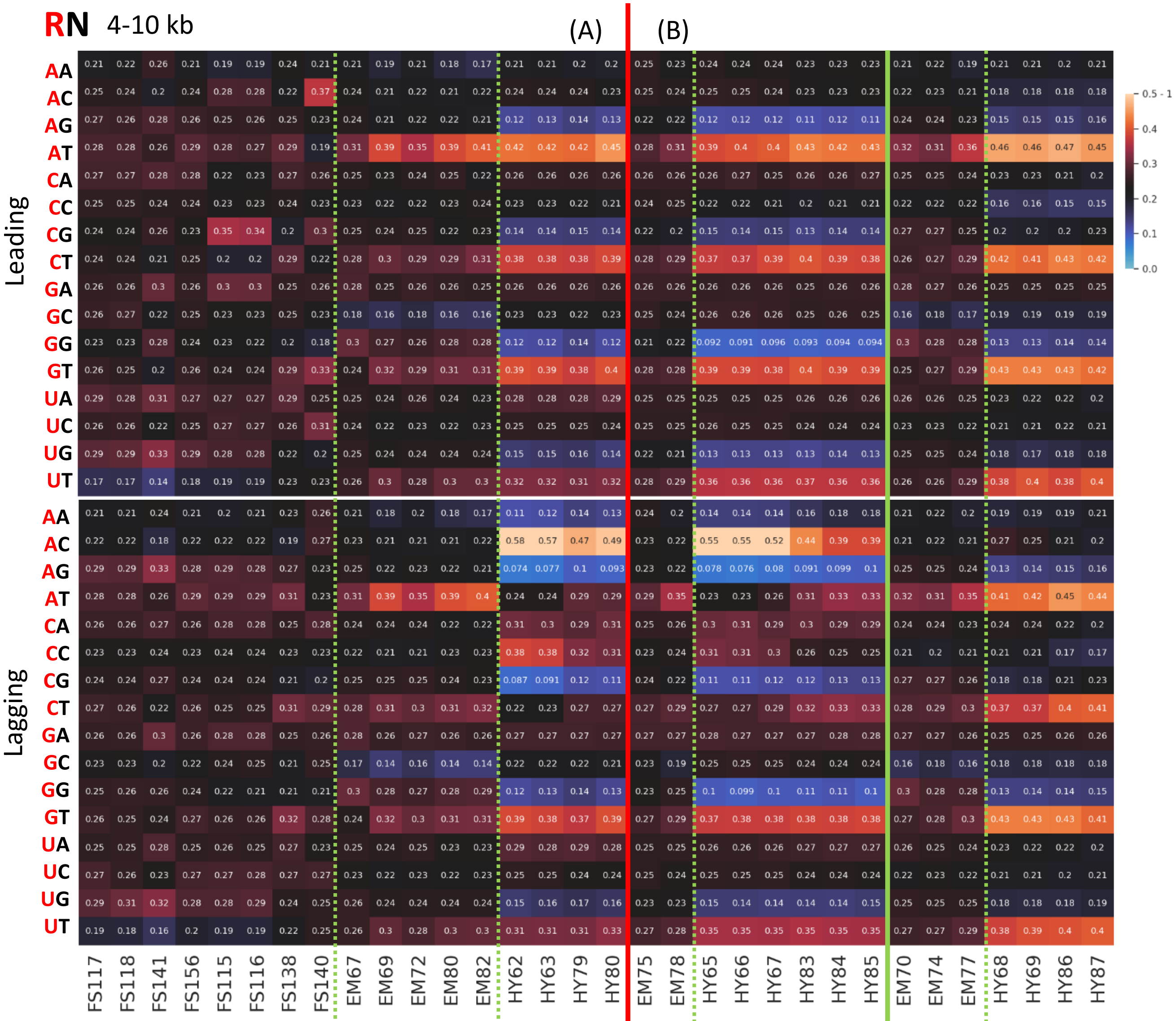


| DNA Pol | WT | | | pol2MG | | pol3LM | pol3LG |
|---|---|---|---|---|---|---|---|
| Method | R | EM | HY | EM | HY | EM | HY |

**Supplementary Figure 12**

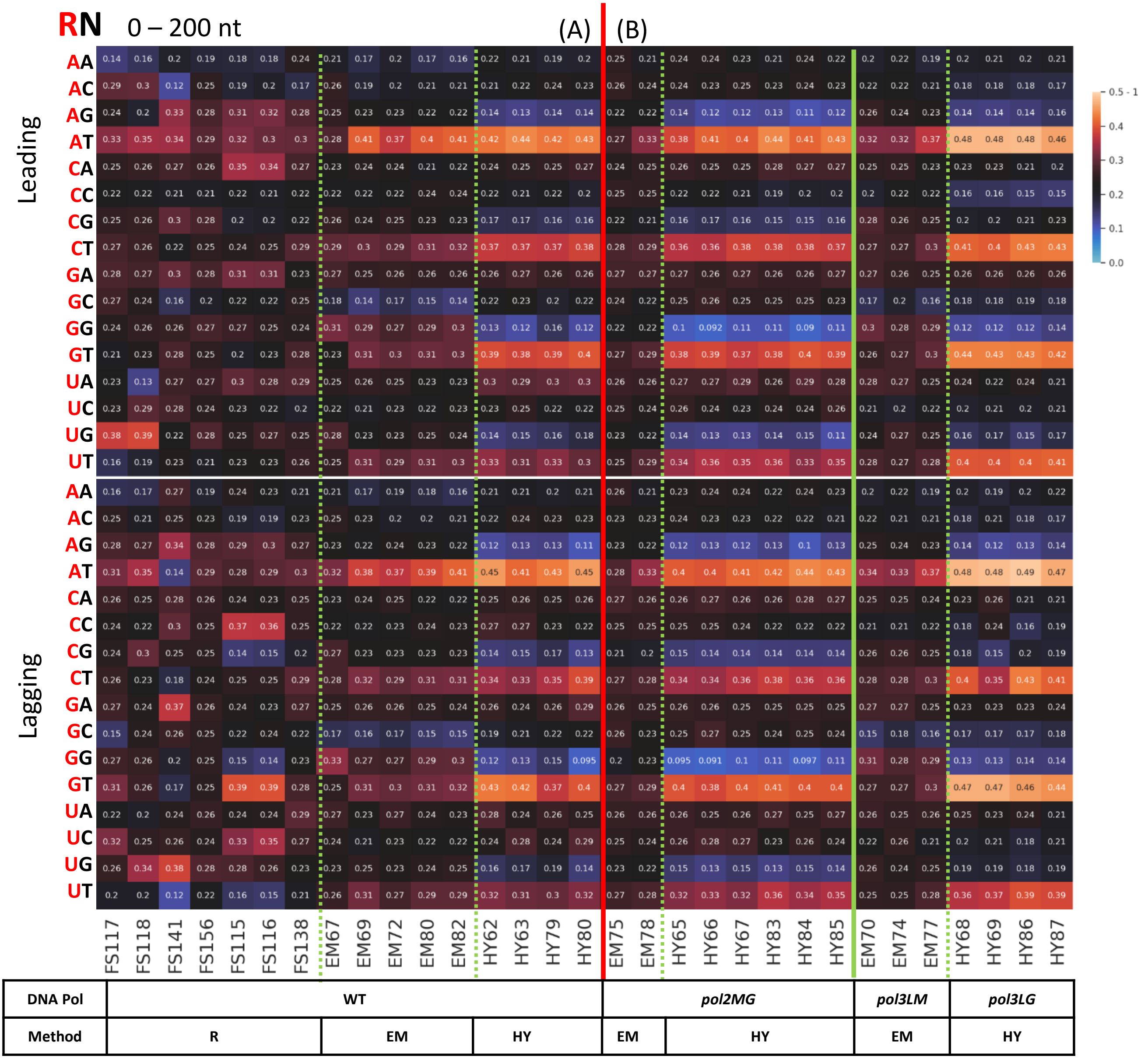


**Supplementary Figure 13**
